# Arginine-dependent hypusination of the eukaryotic translation initiation factor (eIF)5A drives erythroid lineage differentiation

**DOI:** 10.1101/2022.03.21.485053

**Authors:** Pedro Gonzalez-Menendez, Ira Phadke, Meagan E Olive, Kathy L. McGraw, Jessica Platon, Julien Papoin, Hongxia Yan, Marie Daumur, Franciane Paul, Raghavendra G. Mirmira, Axel Joly, Jérémy Galtier, Anupama Narla, Guillaume Cartron, Valérie Dardalhon, Valérie S. Zimmermann, Marc Sitbon, Thomas E. Dever, Narla Mohandas, Lydie Da Costa, Namrata D. Udeshi, Lionel Blanc, Sandrina Kinet, Naomi Taylor

## Abstract

Metabolic programs contribute to hematopoietic stem and progenitor cell (HSPC) fate but it is not known whether the metabolic regulation of protein synthesis controls HSPC differentiation. We discovered that SLC7A1/CAT1-dependent arginine uptake and its catabolism to spermidine control the erythroid specification of HSPCs via activation of eukaryotic translation initiation factor 5A (eIF5A). eIF5A activity is dependent on the metabolism of spermidine to hypusine and inhibiting hypusine synthesis abrogates erythropoiesis and diverts EPO-stimulated HSPCs to a myeloid fate. Proteomic profiling reveals mitochondrial translation to be a critical target of hypusinated eIF5A and induction of mitochondrial function partially rescues erythropoiesis in the absence of hypusine. Within the hypusine network, ribosomal proteins are highly enriched and we identify defective eIF5A hypusination in erythroid pathologies caused by abnormal ribosome biogenesis. Thus, eIF5A-dependent protein synthesis is critical in the branching of erythro-myeloid differentiation and attenuated eIF5A activity characterizes ribosomal protein-linked disorders of ineffective erythropoiesis.

## INTRODUCTION

Hematopoietic stem cells (HSCs) possess two fundamental characteristics, the capacity for self-renewal and the sustained production of all blood cell lineages. The differentiation of HSCs to the erythroid lineage is distinctive in that progressive mitoses result in the production of daughter cells that differ, morphologically and functionally, from their parent cell, with terminal erythroid differentiation generating enucleated reticulocytes. This maturation of erythroid precursors is tightly regulated; each stage of terminal erythroid differentiation is defined by a decreased cell size, chromatin condensation, progressive hemoglobinization, membrane alterations, and changes in gene expression (An et al., 2014; Hu et al., 2013; Li et al., 2014; Ludwig et al., 2019; Schulz et al., 2019). The proper progression of these steps is regulated by transcription factors, cell-cell contacts, oxygen sensors, and cytokines––most notably, erythropoietin. The finding that defects in any of these pathways results in pathological erythroid differentiation underscores their importance in this process (Valent et al., 2018).

Importantly, erythropoiesis is also a formidable process at a metabolic level. The production of 200×10^9^ red cells per day in the bone marrow, and their release into the peripheral circulation, is highly dependent on iron, glucose, fatty acid and amino acid metabolism. Amino acids contribute to multiple aspects of cell metabolism, ranging from precursors of nucleic acids, conversion to glucose and/or lipids, stimulation of the mTOR signaling pathway, production of TCA cycle intermediates, and maintenance of intracellular redox, amongst others (Endicott et al., 2021; Spinelli and Haigis, 2018; Wei et al., 2020). In the context of red cell differentiation, glutamine-dependent nucleotide biosynthesis is a requirement for erythroid commitment (Oburoglu et al., 2016; Oburoglu et al., 2014). Furthermore, glutamine-derived production of succinyl-CoA is necessary for the production of heme (Burch et al., 2018), glutamine- and lysine-induced mTOR signaling promote erythroid maturation (Chung et al., 2015; Gonzalez-Menendez et al., 2021), and glutamine metabolism contributes to oxidative phosphorylation and nucleotide biosynthesis required for erythroid commitment (Chung et al., 2015; Gonzalez-Menendez et al., 2021). Finally, amino acids are also critical for regulating lipid metabolism in erythroid precursors (Huang et al., 2018).

Arginine––a semi-essential dibasic, cationic amino acid––is one of the most versatile amino acids, contributing to the synthesis of urea, nitric oxide, proline, creatine, agmatine, and polyamines. On the basis of these different properties, arginine plays a role in a myriad of physiological and pathological processes including immune function, insulin sensitivity, wound healing, hormone secretion, endothelial function, and cancer proliferation (Morris, 2006). Moreover, arginine has been found to impact erythroid differentiation as well as expression of the SCL7A1/CAT1 arginine transporter (Rotoli et al., 2009; Shima et al., 2006). Notably though, our understanding of the function of arginine metabolism in erythroid development is limited.

Here, we show that SLC7A1/CAT1-mediated arginine uptake and its catabolism to the polyamine spermidine control erythroid commitment and differentiation, respectively. The generation of spermidine is required for hypusination of the eukaryotic translation factor 5A (eIF5A), a post-translational modification resulting from the conjugation of the aminobutyl moiety of spermidine to lysine. Following hypusination, eIF5A promotes translation elongation as well as translation termination (Dever et al., 2014; Dever and Ivanov, 2018; Park and Wolff, 2018). Our quantitative proteomics analyses identified mitochondrial translation as a major target of eIF5A function in erythroid progenitors. This impacted pathway was critical for erythroid differentiation as interventions that augmented mitochondrial metabolism partially rescued erythropoiesis under hypusine-attenuated conditions. Furthermore, genetic alterations in ribosomal proteins (RPs), key components of protein synthesis, drive anemia in bone marrow failure disorders, including Diamond-Blackfan anemia and myelodysplastic syndromes with chromosome 5q deletions. The significant impact of RP mutations on the erythroid lineage highlights the critical nature of global protein synthesis in erythropoiesis (Bello et al., 2018; Khajuria et al., 2018). Given the importance of arginine metabolism on eIF5A-stimulated protein synthesis in erythroid differentiation, we evaluated the impact of RP haploinsufficiency on hypusination. Strikingly, our data identify a novel link between RPs and hypusination in the regulation of erythropoiesis––haploinsufficiency of RPs significantly decreased eIF5A hypusination in erythroid progenitors. Thus, our study reveals a mechanism via which ribosomal proteins regulate hypusination, thereby governing erythroid differentiation.

## RESULTS

### Erythropoietin-stimulated human progenitors are skewed to a myeloid cell fate in the absence of SLC7A1-mediated arginine transport

The availability of intracellular arginine is regulated in large part by cell surface expression of arginine transporters in the SLC7 cationic amino acid transporter (CAT) family (Deves and Boyd, 1998; Verrey et al., 2004) and we found that erythropoietin stimulation of human CD34^+^ HSPCs (**Fig. S1A** and **S1B**) results in a significant upregulation in the cell surface expression of SLC7A1 (p<0.05, **Fig. 1A**). Moreover, this upregulation was associated with a 2-3-fold increase in arginine uptake within 2 days of EPO stimulation (p<0.05, **Figs. 1B** and p<0.01, **S1C**). It is interesting to note that this increase in arginine transport is specific and does not represent a global augmentation in amino acid uptake; rEPO stimulation did not augment transport of glutamine (**Fig. S1C**), an amino acid critical for erythroid development (Oburoglu et al., 2016; Oburoglu et al., 2014). Cell surface SLC7A1 levels then decreased during terminal erythroid differentiation (p<0.001, **Fig. 1C**), associated with a 5-fold reduction in arginine uptake between days 4 and 10 of differentiation (p<0.01, **Fig. 1D**). Arginine was transported into these cells by SLC7A1/CAT1 rather than by the y+LAT/4F2hc heterodimer (Closs et al., 2006), as transport was inhibited by N-ethylmaleimide (NEM), a pharmacological agent that blocks CAT1 but not y+LAT (Beyer et al., 2013). NEM decreased arginine uptake by 9-fold, to basal levels detected on mature human RBCs (p<0.0001, **Fig. S1D**).

**Figure 1.**
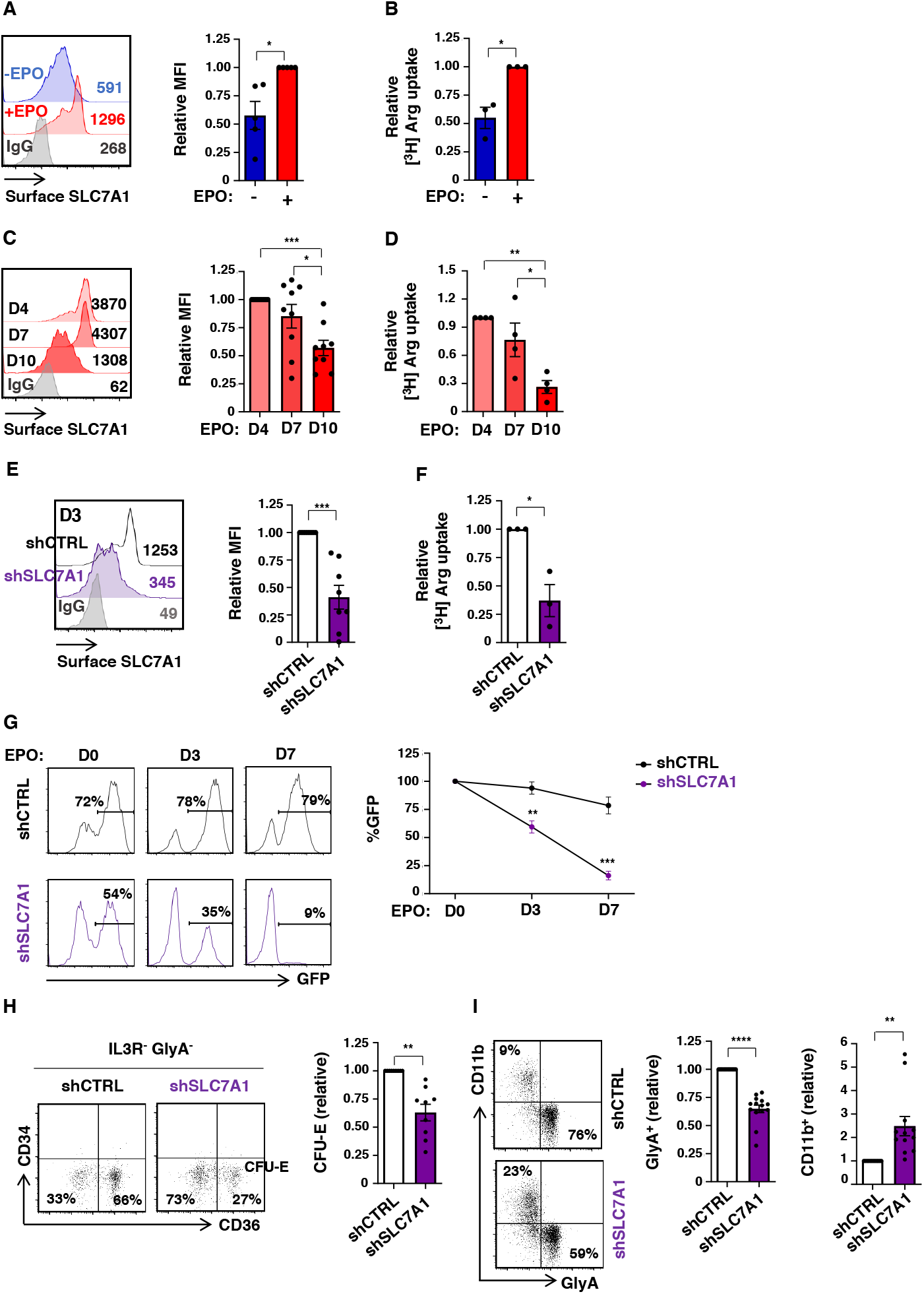
HSC specification to the erythroid lineage is dependent on the expression and function of the SLC7A1 arginine transporter. (**A**) Cell surface expression of SLC7A1 was evaluated following rEPO-induced erythroid differentiation of CD34^+^ progenitors (day 4) and representative histograms in the absence or presence of rEPO are presented (left). Relative MFIs of SCL7A1 in rEPO-induced progenitors are compared with levels detected in the absence of EPO (right; n=5 independent experiments). (**B**) Arginine uptake in the absence or presence of rEPO (day 4) was monitored using L-[2,3,4-^3^H)] arginine monohydrochloride (2 µCi) for 10 min at RT. Uptake in the presence of EPO was arbitrarily set at 1 (means of triplicates in 3 independent experiments). (**C**) SLC7A1 levels were monitored at days 4, 7 and 10 of rEPO-mediated differentiation and representative histograms are shown (left). MFIs of SLC7A1 staining relative to day 4 were quantified (n=9, right). (**D**) Arginine uptake was evaluated at days 4, 7 and 10 of erythroid differentiation; arginine uptake at day 4 was arbitrarily set at 1 (means of triplicates in 4 independent experiments). (**E**) CD34^+^ progenitors were transduced 3 days with GFP-tagged shcontrol (shCTRL) and shSLC7A1 lentiviral vectors and representative histograms of cell surface SLC7A1 expression on GFP^+^ cells (left) as well as quantification of SLC7A1 expression relative to control-transduced cells is shown (right, n=8 independent experiments). (**F**) Arginine uptake was monitored in FACS-sorted progenitors transduced with shCTRL and shSLC7A1 vectors at day 4 and uptake levels relative to the shCTRL condition are presented (n=3). (**G**) The evolution of shCTRL- and shSLC7A1-transduced progenitors was monitored as a function of GFP expression at days 0, 3 and 7 of rEPO-induced differentiation and representative histograms are shown (left). Quantification of the percentages of GFP^+^ cells relative to day 0 is presented (n=6, right). (**H**) Differentiation of rEPO-induced shCTRL- and shSLC7A1-transduced progenitors was monitored as a function of CD34 and CD36 expression on IL3R^-^GlyA^-^ cells; CFU-E are defined by an IL3R^-^GlyA^-^CD34^-^CD36^+^ phenotype (day 3, left). Quantification of CFU-E relative to shCTRL-transduced progenitors is presented (n=9, right). (**I**) The differentiation of progenitors to a erythroid (GlyA) versus myeloid (CD11b) fate was evaluated by flow cytometry at day 3 of rEPO-induced differentiation. Representative plots (left) and quantifications are presented. Surface expression of GlyA and CD11b was monitored on shCTRL- and shSLC7A1-transduced progenitors at day 3 of differentiation (left). Quantification of erythroid (n=14) and myeloid (n=12) differentiation is shown in independent experiments (right). *p<0.05; **p<0.01; ***p<0.001; ****p<0.0001

The significant changes that we detected in SLC7A1 expression and arginine uptake during CD34^+^ HSPC differentiation suggested that SLC7A1-mediated arginine metabolism plays a key role in erythropoiesis. We therefore modulated SLC7A1 expression using an shRNA lentiviral vector harboring the GFP reporter as a transgene. SLC7A1 transcripts as well as cell surface expression were significantly decreased as compared to HSPC transduced with a control (shCTRL) vector (p<0.05, **Fig. S1E** and p<0.001, **Fig. 1E)**. While shSLC7A1-transduced progenitors survived, expansion was decreased by 2-fold (p<0.0001, **Fig. S1F**). Moreover, in the context of EPO stimulation, these cells were negatively selected; there was a significant loss of GFP^+^ progenitors during 7 days of erythroid differentiation, decreasing from 54% to 9% in a representative experiment (p<0.001, **Fig. 1G**). Importantly, a critical role for SLC7A1 was specific to erythroid differentiation as progenitors with downregulated SLC7A1 were not negatively selected following expansion in the absence of erythropoietin (**Fig. S1G**). Consistent with an important function for SLC7A1 in erythroid differentiation, its downregulation resulted in the reduction of IL3R^-^GlyA^-^CD34^-^CD36^+^ CFU-E (p<0.01, **Fig. 1H**) and significantly decreased GlyA, CD36, and CD71 erythroid markers (p<0.0001, **Figs. 1I and S1H**). Even more importantly, these EPO-signaled HSPCs underwent an altered lineage fate, with a >2-fold increase in the differentiation of CD11b^+^ myeloid cells (p<0.01, **Fig. 1I**). Thus, SLC7A1 plays a non-redundant role in regulating the erythroid differentiation of human progenitor cells and in its absence, HSPCs are skewed to a myelomonocytic cell fate.

### Arginine metabolism is required for erythroid lineage commitment as well as terminal erythroid differentiation

To specifically evaluate the role of arginine in the commitment of HSPCs to an erythroid lineage fate, progenitors were stimulated with rEPO in media depleted of arginine. In arginine-depleted media, a significantly lower percentage of progenitors adopted a CFU-E fate (p<0.0001, **Fig. 2A**) and these cells exhibited a significantly decreased proliferation capacity (p<0.05, **Fig. S2A)**. Moreover, under conditions of arginine depletion, sorted CD34^+^CD36^-^ burst forming units-erythroid (BFU-E, **Fig. S2B**, left) exhibited a reduced capacity to differentiate towards a CD34^-^CD36^+^ colony forming unit-erythroid (CFU-E) stage (**Fig. S2B**, right panels). In agreement with the SLC7A1 knockdown experiments (**Fig. 1**), arginine depletion massively skewed progenitors to a CD11b^+^ myeloid fate with an almost complete absence of progenitors expressing GlyA, CD36 or CD71 at both days 3 and 7 of rEPO stimulation (p<0.05, **Figs. 2B and** p<0.0001, **S2C**). Together, these data highlight the critical role of arginine in promoting the commitment of progenitors to an erythroid fate.

**Figure 2.**
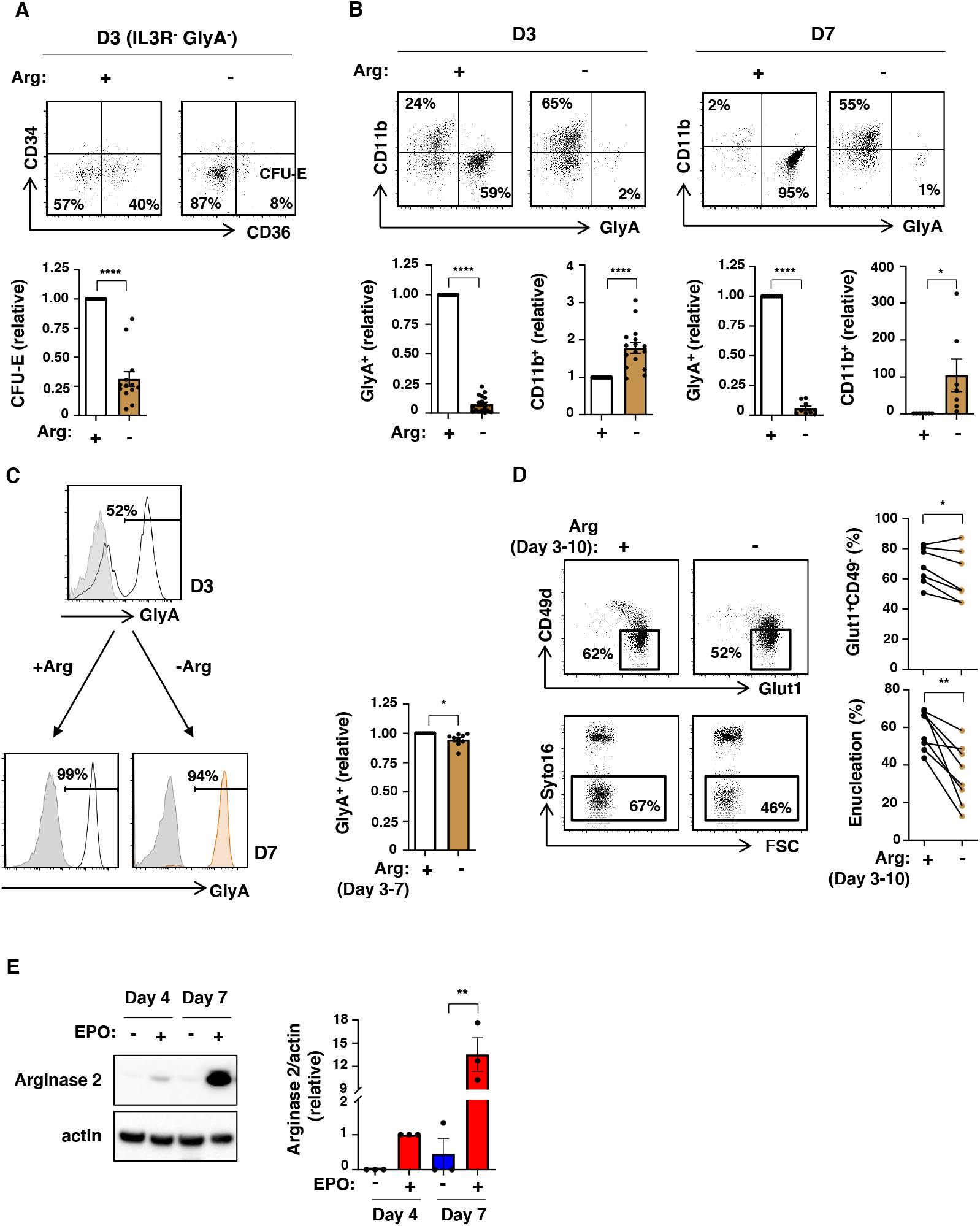
Arginine is required for erythroid lineage commitment and its absence attenuates terminal differentiation. (**A**) The fate of progenitors treated with rEPO in the presence (+) or absence (-) or exogenous arginine was monitored as a function of CD34 and CD36 expression in IL3R^-^GlyA^-^ cells at day 3 of differentiation. Representative dot plots are presented (top) and IL3R^-^GlyA^-^CD34^-^CD36^+^ (CFU-E) cells were quantified relative to differentiation in the presence of arginine (n=13, bottom). (**B**) Relative levels of erythroid and myeloid differentiation were monitored by GlyA and CD11b staining, respectively. Representative dot plots are shown at days 3 and 7 of differentiation in the presence or absence of arginine (top). Quantification of GlyA^+^ (n=17, day 3; n=9, day 7) and CD11b^+^ cells (n=16, day 3; n=7, day 7) are presented (bottom). (**C**) The impact of arginine at later stages of erythroid differentiation was evaluated by depleting arginine following 3 days of EPO-induced erythroid differentiation. GlyA was evaluated at day 3 (top histogram; grey histograms, isotype control; solid line histograms, specific staining) and then 4 days later (day 7), in the presence or absence of arginine (bottom histograms, solid line and orange histograms, respectively). Quantification of the percentages of GlyA^+^ cells was compared following EPO-induced differentiation from day 3-7 in the presence or absence of arginine (n= 9, right). (**D**) The impact of arginine deprivation between days 3 and 10 of rEPO-induced differentiation was monitored as a function of CD49d/GLUT1 profiles (top) and enucleation (Syto16 staining, bottom). Representative plots are shown (left) and quantification in 8 individual donors is presented (right). (**E**) Arginase 2 expression was evaluated by immunoblot at days 4 and 7 of differentiation in the absence or presence of EPO. Actin levels are shown as a loading control (left). Quantification of arginase 2 expression relative to actin was evaluated in the different conditions (right, n=3). *p<0.05; **p<0.01; ****p<0.0001

As SLC7A1 levels decreased during terminal erythropoiesis (**Fig. 1C**), it was of interest to determine whether arginine is required during this stage of differentiation. When arginine was depleted from erythroblasts at day 3 of differentiation, a time point at which GlyA was expressed on approximately 50% of precursors, all cells upregulated GlyA by day 7 (p<0.05, **Fig. 2C**), albeit with a decrease in cell expansion (p<0.001, **Fig. S2D**). Notably though, terminal differentiation as well as enucleation was markedly inhibited, with the latter decreasing from a mean of 59 to 35% (p<0.01, **Fig. 2D**). These data were somewhat surprising given the loss in cell surface SLC7A1 levels upon terminal differentiation (**Fig. 1C**). As such, we evaluated the possibility that intracellular arginine was regulated at the level of arginase expression. Importantly, arginase 2 (ARG2) was upregulated by 12-fold between days 4 and 7 of erythroid differentiation and remained elevated in late-stage erythroblasts (day 14, p<0.01, **Figs. 2E**) while arginase 1 (ARG1), initially expressed at much lower levels, was only significantly upregulated at very late-stage terminal differentiation (day 14, p<0.05, **Fig. S2E**). Thus, the strong induction of ARG2 may be associated with the continued requirement for extracellular arginine throughout the erythroid differentiation process.

### Erythroid lineage specification is dependent on arginine-dependent polyamine biosynthesis

Arginine is a precursor of numerous metabolic intermediates which are generated through distinct catabolic pathways (**Fig. S2F**). To identify which pathway(s) and intermediates are involved in arginine-regulated erythroid differentiation, we first monitored the importance of the arginine-derived synthesis of creatine and nitric oxide. We found that neither of these pathways was required because induction of GlyA was not abrogated by either cyclocreatine or asymmetric dimethylarginine (ADMA), which inhibit creatine and NO production, respectively (**Fig. S2G and S2H**). We next evaluated the role of arginine as a precursor of polyamines **(Fig. 3A)**. Notably, polyamine biosynthesis was found to be strictly required for erythroid differentiation. Inhibition of polyamine biosynthesis via the targeting of ODC (ornithine decarboxylase) or SRM (spermidine synthase) with DFMO (alpha-difluoromethylornithine) and MCHA (trans-4-methyl cyclohexylamine), respectively, dramatically abrogated erythroid differentiation (p<0.001-0.0001, **Figs. 3B and 3C**). While DFMO decreased the proliferation of these progenitors, there was a profound impact on progression to the CFU-E stage, with a lack of induction of the GlyA, CD36, and CD71 terminal erythroid markers (p<0.0001, **Figs. 3B, S3B-S3C**). In accord with the arginine deprivation experiments, progenitors treated with either DFMO or MCHA exhibited a skewed differentiation to a myeloid cell fate, with up to 80% CD11b^+^ cells (p<0.05-0.01, **Figs. 3B-3C**). Similarly, activation of SAT1 (spermidine/spermine N1-acetyltransferase)-mediated polyamine catabolism by DENS (N1, N11-diethylnorspermine) resulted in decreased GlyA upregulation (p<0.0001) and the induction of the CD11b myeloid marker (p<0.05, **Fig. 3D**). Spermidine, and not spermine, was specifically required for erythropoiesis as inhibiting spermine synthase (SMS) with APCHA (N-(3amino-propyl) cyclohexylamine), did not negatively impact erythroid differentiation (**Fig. 3E**). Furthermore, in contrast with requirement of arginine for both erythroid commitment and terminal erythroid differentiation, polyamine biosynthesis was a prerequisite only for erythroid commitment, but not for terminal erythroid differentiation or enucleation (**Figs. S3D and S3E**).

**Figure 3.**
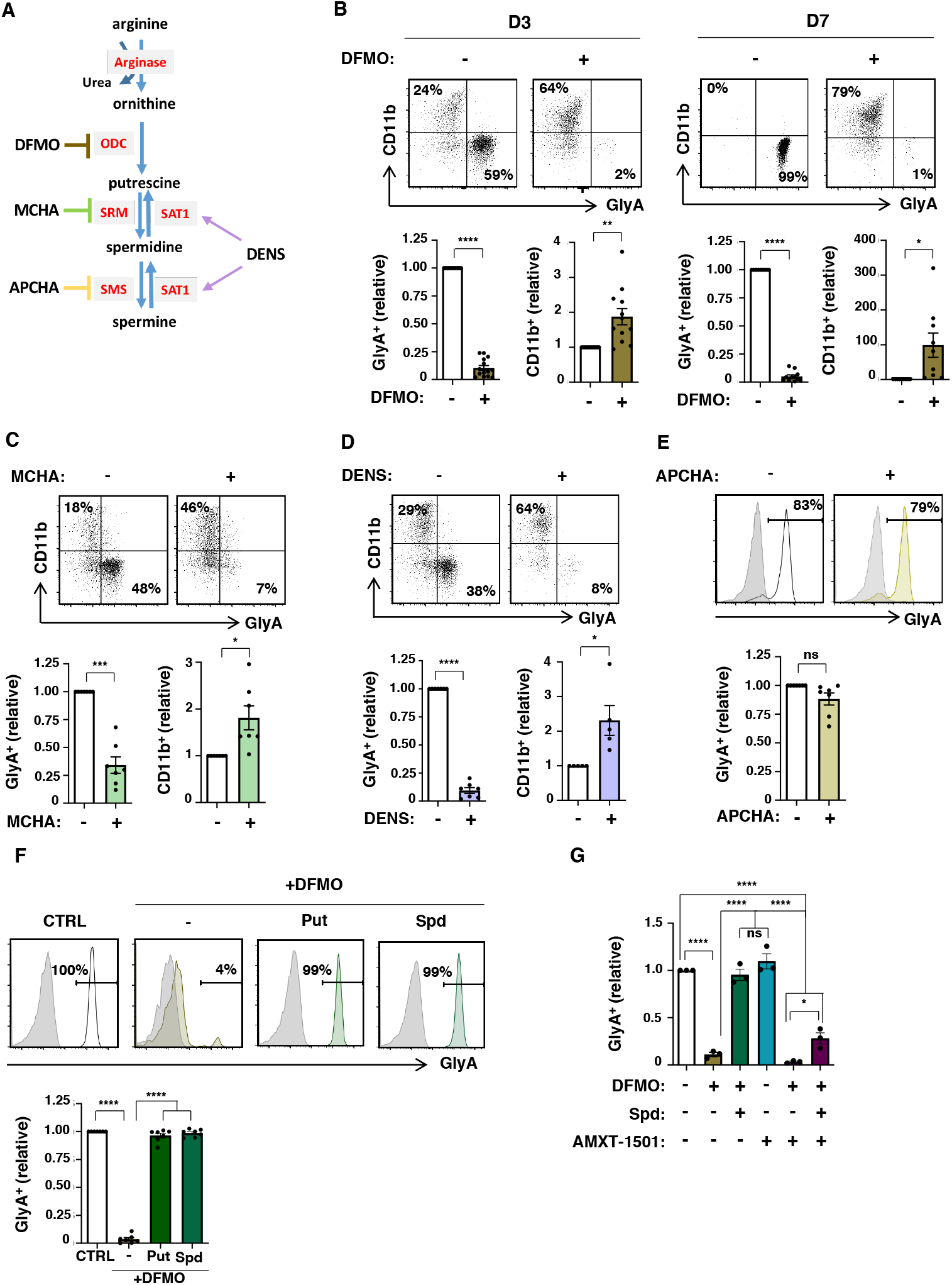
Erythroid differentiation is dependent on arginine-derived polyamine biosynthesis but not transport. (**A**) Schematic representation of polyamine biosynthesis from arginine with key enzymes indicated in red (ODC: ornithine carboxylase, SRM: spermidine synthase, SMS: spermine synthase, SAT1: Spermidine/spermine-N1-acetyl transferase). Steps that are inhibited by DFMO (alpha-difluoromethylornithine), MCHA (trans-4-methyl cyclohexylamine) and APCHA (N-(3amino-propyl) cyclohexylamine) are presented. DENS (N1, N11-diethylnorspermine), an activator of polyamine catabolism, is also indicated. (**B**) The impact of DFMO (1mM) on rEPO-induced differentiation of CD34^+^ progenitors was monitored by evaluating CD11b/GlyA profiles at days 3 and 7. Representative dot plots (top) and quantification of relative levels at days 3 (n=14 for GlyA, n=12 for CD11b) and 7 (n=11 for GlyA, n=9 for CD11b) of differentiation are shown (bottom). (**C**) The impact of MCHA (100µM) was evaluated as a function of CD11b/GlyA profiles (top) at day 3 and quantifications are shown (n=7, bottom). (**D**) The impact of DENS (10µM) on EPO-induced differentiation was evaluated at day 3 and representative plots (top) and quantifications (n=7 for GlyA and n=5 for CD11b) are shown (bottom). (**E**) APCHA (100µM) was added to rEPO-induced progenitors and representative profiles (top) and quantifications (n=7, bottom) are shown. (**F**) DFMO-treated progenitors were differentiated with rEPO in the presence or absence of putrescine (Put, 100µM) or spermidine (Spd, 100µM) and GlyA was evaluated at day 7 (top, green histograms). Quantification from n=7 independent experiments is presented (bottom). (**G**) Erythroid differentiation was induced in the presence or absence of DFMO, spermidine and AMXT-1501 (2.5µM), an inhibitor of polyamine transport. Quantification of GlyA expression relative to control conditions is presented (n=3, bottom) *p<0.05; ***p<0.001; ****p<0.0001; ns, not significant

To assure that the impact of pharmacological inhibitors was directly coupled to polyamine biosynthesis, and not to any off-target effects, we performed rescue experiments and found that ectopic putrescine and spermidine completely restored the DFMO- and MCHA-mediated attenuation of erythroid differentiation (p<0.0001, **Fig. 3F and S3F**). It was also interesting to note that under control conditions, erythroid differentiation was not dependent on polyamine transport from extracellular stores as it was not blocked by the AMXT-1501 polyamine transporter inhibitor (Burns et al., 2009) (**Fig. 3G**). However, following DFMO-mediated abrogation of intracellular polyamine biosynthesis, AMXT-1501 abrogated the potential of spermidine to rescue erythroid differentiation (p<0.0001, **Figs. 3G and S3G**). These data point to an evolutionary redundancy; intracellular polyamine synthesis is sufficient for erythroid differentiation under physiological conditions but under stress conditions wherein intracellular polyamine levels are limiting, the transport of extracellular polyamines can promote erythropoiesis.

### The DHPS-catalyzed generation of hypusine from spermidine is required for erythroid differentiation

Polyamines are also involved in numerous functions that are critical for cell growth and homeostasis. Indeed, their pleiotropic nature has made it difficult to ascribe specific roles to polyamines (Igarashi and Kashiwagi, 2010). Polyamines can directly stimulate protein synthesis, but the polyamine spermidine also serves as a precursor for the generation of the natural amino acid hypusine (N ϵ-4-amino-2-hydroxybutyl(lysine)), an essential post-translational modification of the eukaryotic translation initiation factor 5A (eIF5A). Hypusine is formed by the conjugation of the aminobutyl moiety of spermidine to lysine-50 of human eIF5A (Cano et al., 2008). Remarkably, eIF5A is the only protein in which this modification is known to occur, and its presence is required for its function––promoting translation elongation, especially of a subset of proteins, as well as translation termination (Dever et al., 2014; Dever and Ivanov, 2018; Park and Wolff, 2018). The synthesis of hypusine is catalyzed through sequential enzymatic steps involving deoxyhypusine synthase (DHPS) and deoxyhypusine hydroxylase (DOHH, **Fig. 4A**).

**Figure 4.**
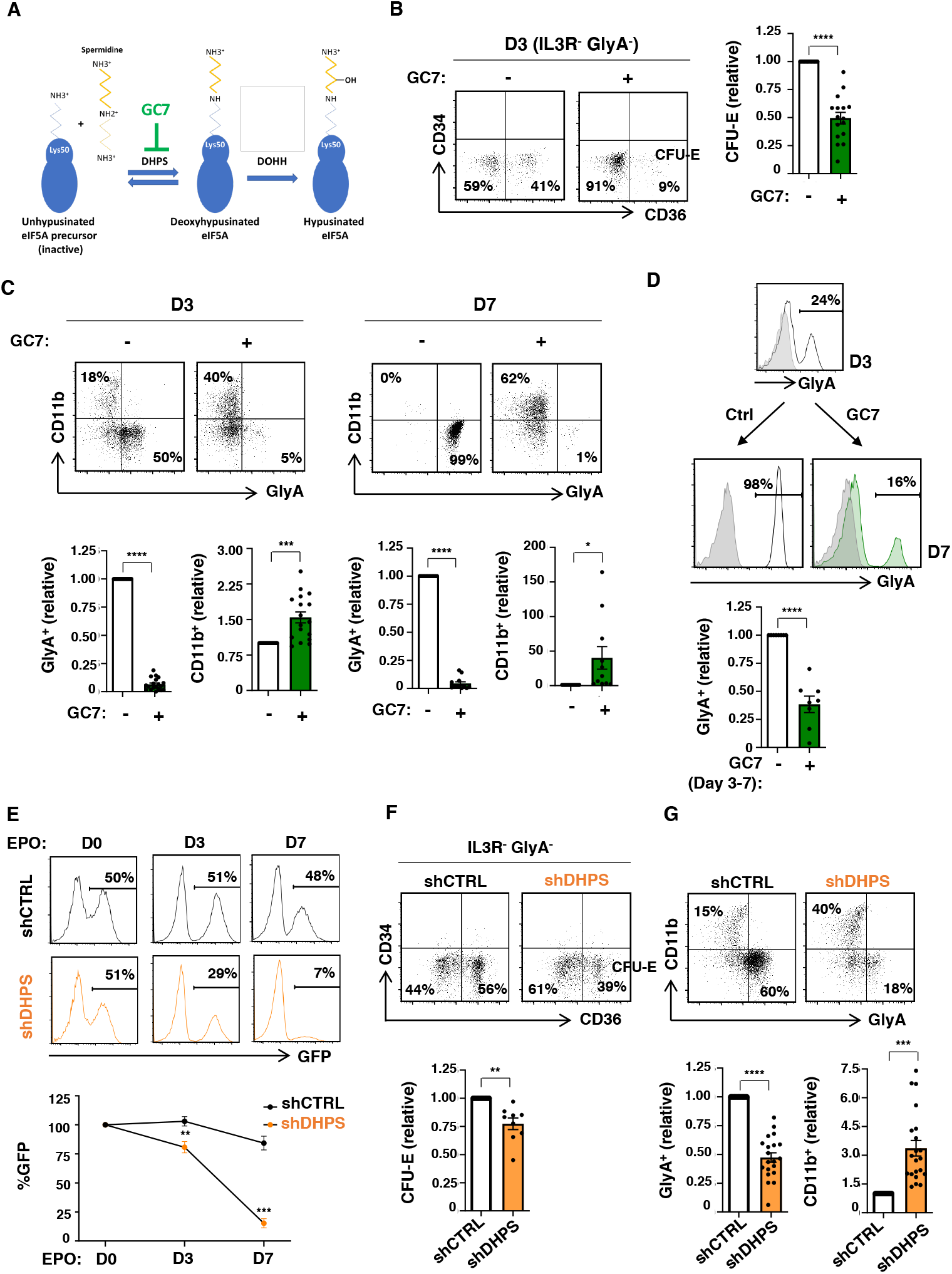
Deoxyhypusine synthase is required for erythroid commitment and differentiation. (**A**) Schematic representation of the two-step process resulting in the hypusination of eukaryotic initiation factor 5A (eIF5A^H^). In the first step, deoxyhypusine synthase (DHPS) catalyzes the addition of an aminobutyl moiety from the spermidine to the lysine 50 on eIF5A, generating an eIF5A intermediate. Subsequently, deoxyhypusine hydroxylase (DOHH) catalyzes the hydroxylation of the spermidine-modification, generating an active hypusinated eIF5A. N1-guanyl-1,7-diamineoheptane (GC7) can inhibit the first step of this reaction. (**B**) The effect of GC7 on early erythropoiesis was evaluated by monitoring CD34/CD36 profiles of EPO-stimulated CD34^+^ progenitors in the absence (-) or presence (+) of GC7 (5µM). Representative dot plots of IL3R^-^GlyA^-^ cells are shown and the percentages of IL3R^-^GlyA^-^CD34^-^CD36^+^ (CFU-E) are indicated (left). Quantification of CFU-E in 16 independent experiments are presented (right). (**C**) Representative dot plots of GlyA/CD11b profiles are presented at days 3 and 7 of EPO-induced differentiation in the absence or presence of GC7 (top). Quantification of GlyA^+^ and CD11b^+^ cells relative to levels in the absence of GC7 (indicated as “1”) are presented (bottom, n=11-19). (**D**) CD34^+^ progenitors were differentiated in rEPO for 3 days and differentiation continued until day 7 in the absence or presence of GC7 (between days 3 and 7). Representative histograms showing GlyA expression at days 3 and 7 (with isotype controls, grey histograms) are presented and quantification of GlyA^+^ cells are presented relative to levels in the absence of GC7 (designated as “1”, bottom, n=8). (**E**) CD34^+^ progenitors were transduced with shControl (CTRL)- or shDHPS lentiviral vectors harboring the eGFP transgene and rEPO added 72h later. GFP expression was monitored at this time point (designated day 0), as well at days 3 and 7 of differentiation and representative are shown (top). Quantification of the evolution of GFP expression relative to day 0 (designated as “100%”) is presented for 5 donors (bottom). (**F**) shCTRL and shDHPS transduced progenitors were differentiated in the presence of rEPO for 3 days and representative CD34/CD36 profiles of IL3R^-^GlyA^-^ cells are presented (top). Quantification of IL3R^-^GlyA^-^CD34^-^CD36^+^ cells in shDHPS-transduced cells relative to shCTRL-transduced cells are shown (bottom, n=9). (**G**) shCTRL and shDHPS transduced progenitors were differentiated for 3 days and representative GlyA/CD11 dot plots are shown (top). Quantification of GlyA^+^ and CD11b^+^ cells are presented relative to shCTRL conditions (bottom; n=21 for GlyA, n=20 for CD11b). *p<0.05; **p<0.01; ***p<0.001; ****p<0.0001

Addition of the spermidine analog N1-guanyl-1,7-diaminoheptane (GC7), one of the most potent inhibitors of DHPS (Jakus et al., 1993) (**Fig. 4A**), significantly blocked the EPO-induced generation of CFU-E (p<0.0001, **Fig. 4B**). Furthermore, myeloid skewing occurred despite a decrease in progenitor expansion in this condition (p<0.0001, **Fig. S4A**). A 30-fold increase in CD11b^+^ cells was detected by day 7 of EPO-induced differentiation (p<0.05, **Fig. 4C**). This increase was accompanied by a massive decrease in erythroid progenitors, irrespective of whether GC7 was added at day 0 or day 3 of EPO-induced differentiation (p<0.0001, **Figs. 4C, 4D, S4B, and S4C**). Upon inhibition of DHPS activity at day 3 of differentiation, the generation of late stage CD49d^-^ orthochromatic erythroblasts was significantly attenuated (p<0.01) and enucleation decreased from a mean of 61 to 13% (p<0.01, **Fig. S4D**). A pharmacological inhibitor of DOHH, the iron chelator ciclopirox olamine (CPX) (Abbruzzese et al., 1991), also significantly inhibited the induction of GlyA to levels <10% of that detected in control conditions (p<0.001, **Fig. S4E** and **S4F**).

To directly evaluate the role of enzymes regulating hypusination, we utilized an shRNA-mediated approach, targeting DHPS. shDHPS-transduced progenitors exhibited a 75% decrease in the DHPS mRNA (p<0.01, **Fig. S4G**), associated with a decreased expansion (p<0.0001, **Fig. S4H**). Moreover, progenitors with downregulated DHPS were massively counter-selected during the first 7 days of erythroid differentiation (p<0.001, **Fig. 4E**). Similar to the impact of GC7, shDHPS-transduced progenitors exhibited a decreased generation of CFU-E (p<0.01, **Fig 4F**) and a significantly attenuated upregulation of GlyA, CD36, and CD71 erythroid markers (p<0.0001, **Fig. 4G and S4J**). Importantly though, the negative impact of DHPS downregulation was restricted to erythroid-committed progenitors; in the absence of rEPO, shDHPS-transduced progenitors were not negatively selected (**Fig. S4I**) and there was a 3-fold expansion of CD11b^+^ myeloid cells (p<0.001, **Fig. 4G**). Thus, in a manner analogous to conditions of decreased arginine uptake, inhibition of DHPS skewed the differentiation of erythroid progenitors to a myeloid cell fate.

### Hypusination of eIF5A in erythroid progenitors regulates the translation of proteins involved in cell cycle, protein metabolism and mitochondrial translation

The data presented above strongly suggested a critical role for hypusination in erythroid differentiation. However, it is notable that eIF5A hypusination (eIF5A^H^) has never been evaluated as a function of erythroid differentiation. Notably, we found that eIF5A levels increased within 4 days of EPO stimulation (p<0.01, **Fig. 5A**). However, hypusine levels decreased during terminal differentiation, with a significant drop between polychromatic and orthochromatic erythroblasts (p<0.05, **Fig. 5B** and p<0.001, **S5A**). Thus, eIF5A hypusination increases during erythroid commitment and then decreases during late-stage terminal differentiation.

**Figure 5.**
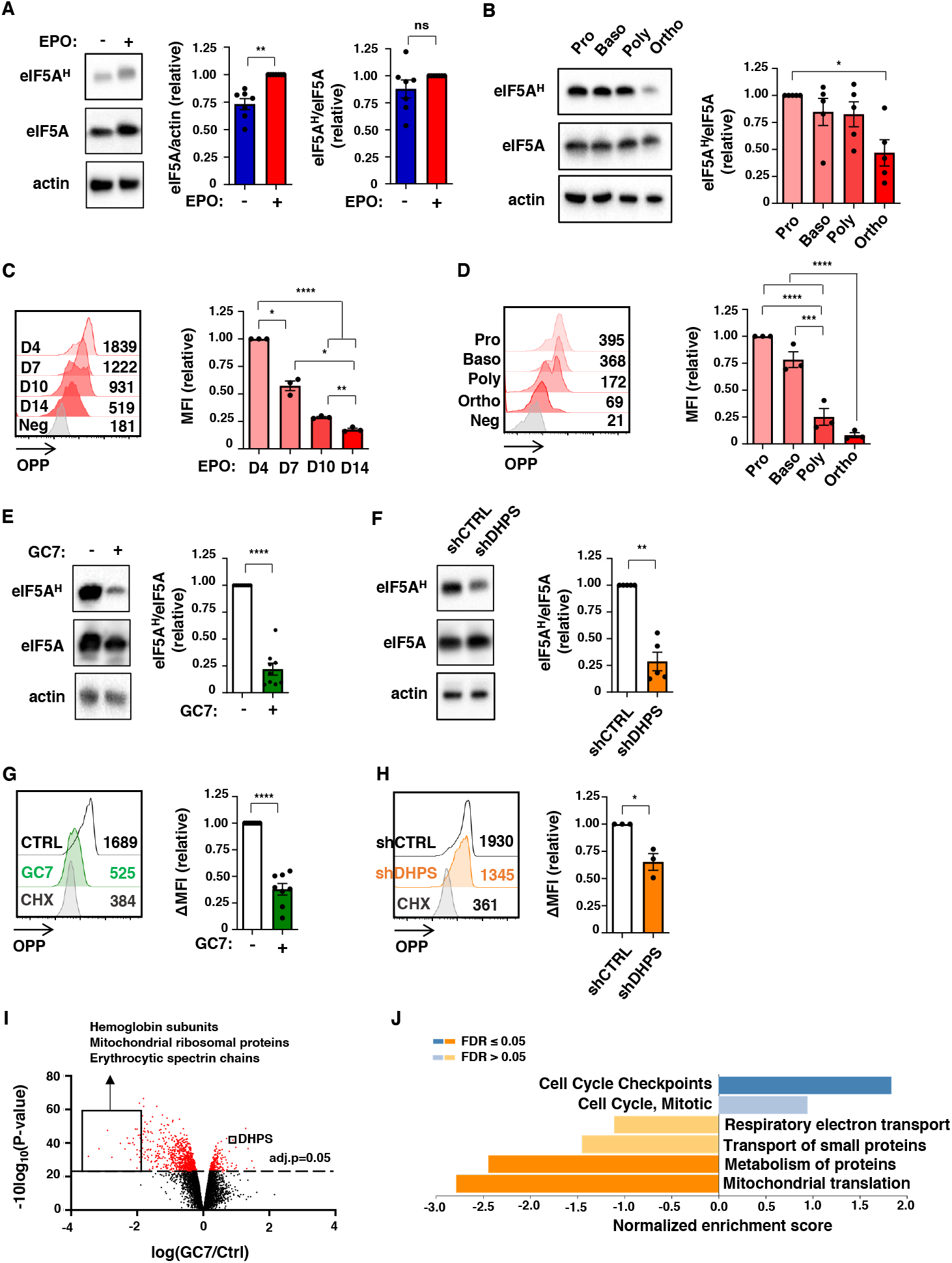
DHPS-mediated hypusination of eIF5A regulates protein translation in early erythroid progenitors. (**A**) Hypusination was evaluated in progenitors differentiated in the presence or absence of EPO (day 4) and representative immunoblots of eIF5A^H^, eIF5A and actin are shown (left). Quantification of eIF5A/actin (middle) and eIF5A^H^ /eIF5A (right) ratios are presented relative to levels in the presence of EPO (n=7). (**B**) Pro-erythroblasts (Pro), basophilic (Baso), polychromatic (Poly) and orthochromatic (Ortho) erythroblasts were sorted on the basis of their GLUT1/CD49d prolife at day 7 of differentiation and hypusination was monitored by immunoblotting (left) and hypusination relative to eIF5A levels was evaluated (n=5, right). (**C**) Protein synthesis was monitored at the indicated day of erythroid differentiation by O-propargyl-puromycin (OPP) labeling and representative histograms and MFI are indicated (left). Quantification of MFIs relative to day 4 are presented for three donors (right). (**D**) Protein synthesis was monitored by OPP labeling in erythroblast subsets 24h following sorting (as in panel B, left). Quantification of MFIs relative to pro-erythroblasts are presented for three donors (right). (**E**) Hypusination was evaluated following 3 days of EPO stimulation in the absence (-) or presence (+) of GC7 (5µM) and representative immunoblots of eIF5A^H^, eIF5A, and actin are shown (left). Quantification relative to levels in the absence of GC7 are presented (n=9, right). (**F**) Hypusination in shCTRL- and shDHPS-transduced progenitors was evaluated at day 3 of differentiation following sorting on the basis of GFP expression and representative plots are shown (left). Quantification of the relative levels of eIF5A^H^/eIF5A is presented (right, n=5). (**G**) CD34^+^ progenitors were differentiated in the presence of EPO, together with GC7 (5µM) or cycloheximide (CHX, 1µM). Protein synthesis was evaluated at day 1 of differentiation and a representative histogram is shown (left). Quantification of protein synthesis relative to control conditions is presented (n=8, right). (**H**) shCTRL and shDHPS-transduced progenitors were sorted 72h post transduction on the basis of GFP expression. Protein synthesis was evaluated 24h following addition of EPO and a representative histogram (left) as well as quantification (right, n=3) are presented. (**I**) CD34^+^ progenitors were differentiated with EPO in the absence or presence of GC7 (5µM) for 2 days and protein expression was evaluated by mass spectrometry-based quantitative proteomics. A volcano plot shows differences in protein expression (Log2FC) induced by GC7 are presented and the identity of specified downregulated and upregulated proteins are noted. Statistical significance of relative protein expression is computed via two-sample moderated T test, and proteins with an FDR-adjusted p<0.05 are colored in red. (**J**) Over-representation analyses of gene ontology (GO) for non-redundant biological processes were evaluated for significantly upregulated and downregulated and enrichment scores are indicated. *p<0.05; **p<0.01; ***p<0.001; ****p<0.0001; ns, not significant

The potential impact of hypusination on protein synthesis was evaluated as a function of the incorporation of the alkyne analog of puromycin, O-propargyl-puromycin (OPP)– –forming covalent conjugates with nascent polypeptide chains that can be evaluated by flow cytometry (Hidalgo San Jose and Signer, 2019; Liu et al., 2012). Within 1 day of EPO stimulation, OPP incorporation was significantly increased, indicating an augmented level of protein synthesis (p<0.05, **Fig. S5B**). Furthermore, OPP staining decreased as a function of erythroid differentiation (**Fig. 5C** and **5D**). Interestingly, the dramatic drop in protein synthesis occurred earlier than the drop in hypusination, between the basophilic and polychromatic erythroblast stages (p<0.001, **Fig. 5D**). These data correlate with previous findings showing that total protein levels also drop between the basophilic and polychromatic stages of differentiation (Gautier et al., 2016).

To assess whether hypusination in hematopoietic progenitors is directly regulated by arginine metabolism, we evaluated hypusination under conditions of arginine deprivation as well as polyamine biosynthesis inhibition. Both these conditions dramatically reduced hypusination, without a significant impact on eIF5A levels themselves (**Figs. S5C and S5D**). Furthermore, polyamine biosynthesis was directly required for hypusination; the DFMO-mediated inhibition of hypusination was rescued by both putrescine and spermidine (p<0.001, **Fig. S5D**). Thus, hypusination in hematopoietic progenitors is strictly dependent on arginine-derived polyamine biosynthesis.

As expected from these results, the inhibition of DHPS––by either GC7 or shRNA-mediated downregulation of the enzyme––resulted in a 75% loss of hypusine as compared to control progenitors (p<0.0001, p<0.01, **Figs. 5E and 5F**). Moreover, these modulations resulted in functional changes in protein synthesis. OPP labelling revealed a significant loss in nascent protein synthesis in both conditions with labeling in GC7-treated progenitors decreasing to levels detected in the presence of cycloheximide, an inhibitor of protein synthesis (p<0.0001, p<0.05, **Figs. 5G and 5H**).

To specifically evaluate the proteins that were regulated by hypusinated eIF5A (eIF5A^H^) in erythroid progenitors, we analyzed progenitors differentiated for 48h in the absence or presence of GC7 by quantitative liquid chromatography tandem mass-spectrometry (LC-MS/MS) based proteomics using tandem mass tags (TMT) (Mertins et al., 2018). Notably, and in agreement with the impact of GC7 on erythroid differentiation, some of the most highly downregulated proteins were hemoglobin subunits and erythroid spectrin chains (**Fig. 5I**). Furthermore, DHPS was significantly upregulated following GC7 treatment, pointing to a potential compensatory feedback loop. Gene set enrichment analysis and protein-protein interaction (PPI) network clustering analysis (**Fig. 5J and Fig. S5E**) revealed a massive loss of proteins involved in mitochondrial translation. This is of interest given previous work from our group and others showing the importance of mitochondrial ROS and protein translation in erythroid differentiation (Gonzalez-Menendez et al., 2021; Liang et al., 2021a; Liu et al., 2017). Furthermore, the specific inhibition of mitochondrial protein synthesis with chloramphenicol resulted in a significant decrease in the generation of GlyA^+^ erythroid progenitors (p<0.001), and a skewing towards the generation of CD11b^+^ myeloid cells (p<0.05, **Fig. S5F**). Thus, while there is a large literature showing the specific importance of protein synthesis during erythropoiesis (Iskander et al., 2021; Khajuria et al., 2018; Vatikioti et al., 2019), our results highlight the importance of the cytoplasmic hypusine-dependent synthesis of the mitochondrial translation apparatus and thus, mitochondrial translation in erythroid differentiation.

### The attenuated mitochondrial metabolism and erythroid differentiation associated with defective hypusination are partially rescued by succinate

To decipher the impact of hypusination on the synthesis of proteins involved in mitochondrial metabolism, we first evaluated expression of mitochondrial complex enzymes. Interestingly, complex II, III and V were all upregulated by EPO-induced differentiation (p<0.05, **Fig. S6A**). Notably though, both GC7 and DHPS downregulation resulted in decreased expression of all 5 mitochondrial complex enzymes while actin levels remained stable (**Fig. 6A**). These data unveil the importance of hypusination in the expression levels of this subset of mitochondrial proteins.

**Figure 6.**
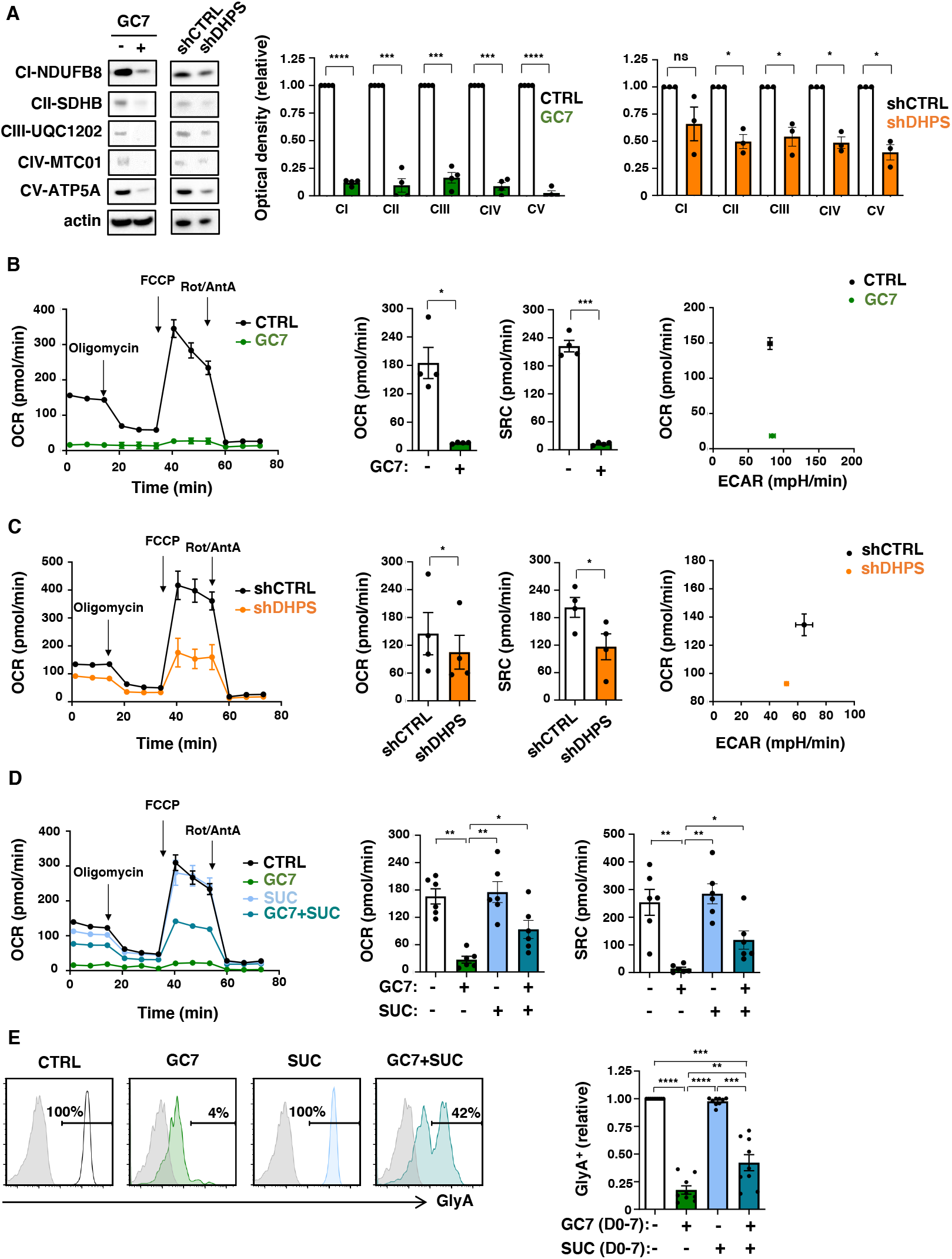
Hypusination-induced oxidative phosphorylation is required for the erythroid commitment of hematopoietic progenitors. (**A**) Mitochondrial complexes (CI to CV) were monitored on progenitors treated with GC7 or following transduction with shCTRL or shDHPS vectors (day 3) using the OXPHOS monoclonal antibody cocktail (left). Quantification relative to control conditions was determined (n=4 for GC7 and n=3 for shRNA-transduced progenitors, middle and right). (**B**) Oxygen consumption rate (OCR), a measure of OXPHOS, was monitored on day 1 of erythroid differentiation in the absence or presence of GC7 (5µM) on a Seahorse XFe96 analyzer following sequential injection of oligomycin, FCCP and Rotenone/Antimycin A (arrows; left panel). Mean basal OCR and SRC levels ± SEM are presented (n=4, middle panels). Representative energy plots of basal OCR and extracellular acidification rate (ECAR), a measure of glycolysis, are presented (right). (**C**) OCR was monitored on FACS-sorted shCTRL- and shDHPS-transduced progenitors at day 1 of differentiation (left). Basal OCR and SRC levels ± SEM were evaluated in 4 independent experiments (middle panels) and a representative OCR/ECAR energy plot is presented (right). (**D**) OCR was monitored on CD34^+^ progenitors differentiated with EPO for 24h in the absence or presence of GC7 (5µM) and succinate (SUC, 5mM) and representative graphs are shown (left). Basal OCR and SRC levels in 6 independent experiments are presented (right). (**E**) Erythroid differentiation in progenitors treated with EPO in the absence or presence of GC7 or succinate was evaluated at day 7 as a function of GlyA expression and representative histograms are shown (right). Quantification of the percentages of GlyA^+^ cells relative to control conditions are presented (n=9). *p<0.05; **p<0.01; ***p<0.001, ****p<0.0001

Based on the importance of eIF5A^H^ for the synthesis of mitochondrial proteins, we studied mitochondrial metabolism in these cells. While GC7 treatment and DHPS downregulation both resulted in diminished mitochondrial biomass and polarization potential (p<0.01, **Fig. S6B** and p<0.05, **S6C**), decreases were less than 1.2-fold. Notably though, the impact on oxidative phosphorylation was much more striking; GC7 almost completely abrogated the capacity of these progenitors to undergo OXPHOS and while the cells exhibited a higher glycolytic capacity, their glycolytic reserve was decreased (**Figs. 6B** and **S6D**). Similarly, inhibition of polyamine biosynthesis with either DFMO or DENS decreased basal respiration and spare respiratory capacity (SRC) by 2.5-4.5-fold (**Fig. S6E**). This loss of OXPHOS, albeit with a maintained glycolysis, was maintained in progenitors with downregulated DHPS (**Fig. 6C** and **S6F**), again highlighting the critical role of hypusination for mitochondrial metabolism. As a control, we used ciclopirox, an inhibitor of hypusination and mitochondrial respiration (Qi et al., 2020), and found that this treatment also resulted in a significantly lower basal and maximal oxygen consumption (p<0.05, p<0.001, **Fig. S6G**).

The decreased level of mitochondrial protein synthesis under conditions where hypusination and mitochondrial protein translation were negatively impacted led us to test the hypothesis that drivers of the TCA cycle would augment erythroid differentiation. Succinate, by bypassing complex I, can potentially provide a fuel source for compromised mitochondria (Phang et al., 2016). Furthermore, our protein analyses revealed a significant loss of the complex II SDHB enzyme (**Fig. 6A**), thereby attenuating the generation of succinate. Of note, addition of succinate to GC7-treated EPO-induced progenitors resulted in a significant upregulation of OXPHOS. The level of both basal and maximal OXPHOS was lower than that detected in control conditions but oxygen consumption was significantly higher than that detected in the presence of GC7 alone (p<0.01, **Fig. 6D**). Most importantly though, the addition of succinate promoted the erythroid lineage commitment of GC7-treated hematopoietic progenitors (p<0.01, **Fig. 6E**). Together, these data reveal an interplay between hypusination and mitochondrial metabolism in regulating erythroid lineage commitment and differentiation.

### Defective erythroid differentiation in ribosomal protein haploinsufficiency is coupled to an attenuated hypusination

Within the hypusine network, Sievert and colleagues found that the most highly enriched eIF5A-interacting proteins are associated with ribosomal function, including RPL and RPS ribosomal proteins (Sievert et al., 2012). Of note, mutations resulting in impaired ribosome biogenesis, including allelic variations in more than 20 different ribosomal proteins (RPs) have all been found to result in Diamond-Blackfan Anemia (DBA). This rare congenital disease predominantly affects erythroid lineage cells and is characterized by macrocytic anemia and bone marrow failure (Da Costa et al., 2020). Furthermore, haploinsufficiency of the *RPS14* gene results in the macrocytic anemia in patients with del(5q) myelodysplastic syndrome (MDS) (Ebert et al., 2008). However, to date, there has been no evaluation of a potential association between RP function and hypusination in erythroid differentiation. We therefore first assessed hypusination in immortalized haploinsufficient *Rps14*^*+/-*^*Mx1Cre*^*+*^ (Schneider et al., 2016) as compared to control *Mx1Cre*^*+*^ murine hematopoietic progenitors. Notably, eIF5A^**H**^ was significantly lower in the former while expression of eIF5A was similar in *Rps14*^*+/-*^*Mx1Cre*^*+*^ and *Mx1Cre*^*+*^ progenitors (p<0.01, **Fig. 7A**). Moreover, similar to human hematopoietic progenitors where an attenuated hypusination was associated with a diminished OXPHOS (**Fig. 6**), basal oxygen consumption as well as spare respiratory capacity were significantly lower in *Rps14*^*+/-*^*Mx1Cre*^*+*^ than *Mx1Cre*^*+*^ progenitors (p<0.01, **Fig. 7B** and **S7A**), thus linking RPS14 expression to hypusination and mitochondrial metabolism.

**Figure 7.**
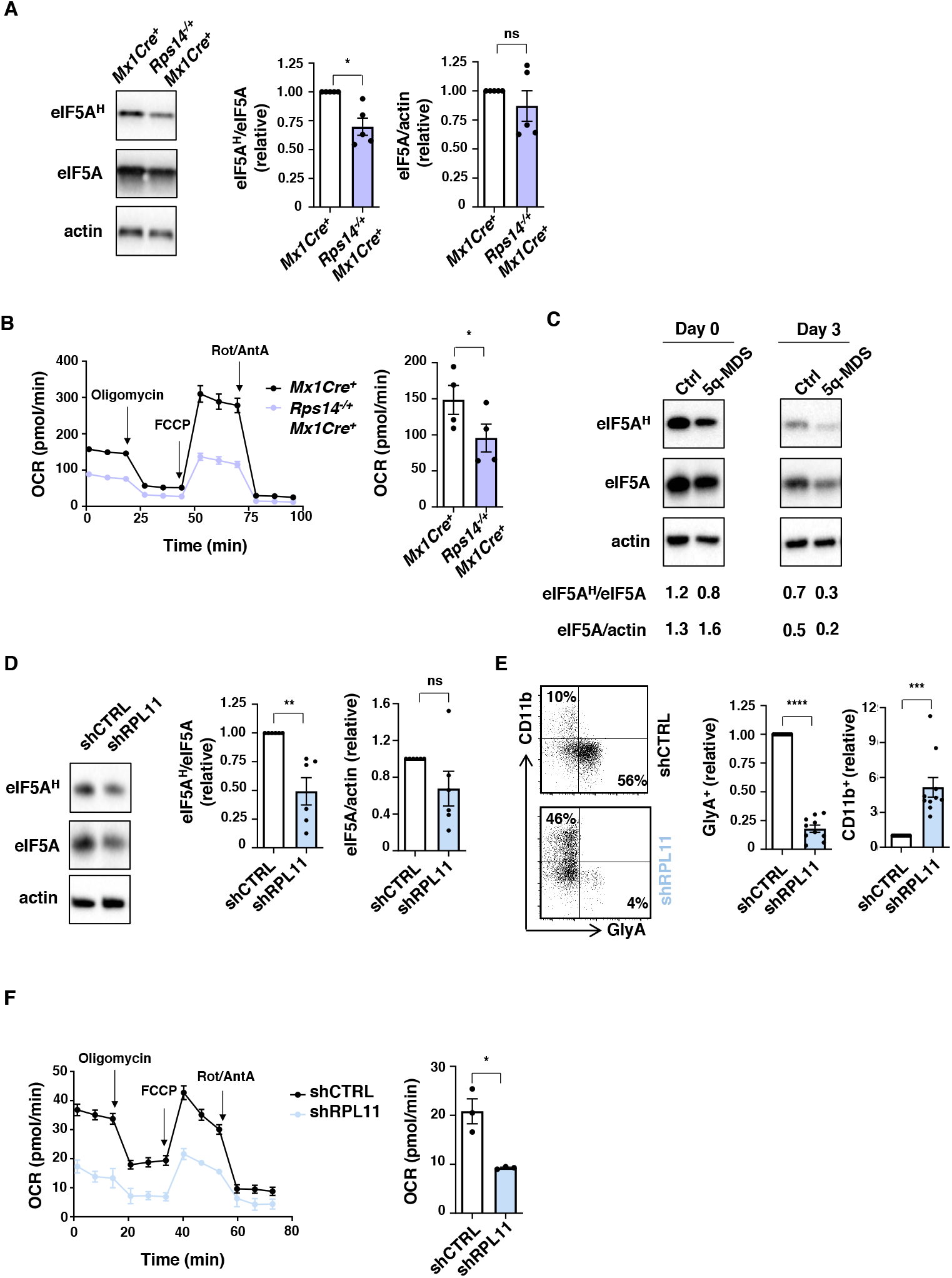
Haploinsufficiency in ribosomal protein genes is coupled to an attenuated hypusination in erythroid progenitors. (**A**) Hypusination and total eIF5A levels were evaluated in immortalized erythroid bone marrow progenitors generated from control *Mx1Cre*^*+*^ and *RPS14* haploinsufficient *Rps14*^*+/-*^*Mx1Cre*^*+*^ mice (Schneider et al., 2016). Representative immunoblots (left) and quantifications of eIF5A^H^/eIF5A and eIF5A/actin levels are presented (right, n=5). (**B**) OCRs of immortalized *Mx1Cre*^*+*^ and *Rps14*^*+/-*^*Mx1Cre*^*+*^ progenitors were evaluated and representative data (left, n=6 technical replicates) (n=4 independent experiments, right panel) are presented. (**C**) CD34^+^ BM progenitors from a healthy control and a del(5q)-myelodysplastic syndrome patient were expanded *ex vivo* and evaluated for hypusination and total eIF5A levels at day 0 and 3 of EPO-induced differentiation. Immunoblots (top) and quantifications (bottom) are presented. (**D**) Hypusination was evaluated in human progenitors transduced with shCTRL and shRPS11 vectors at day 3 of differentiation. Representative immunoblots of eIF5A^H^ and total eIF5A levels in shCTRL- and shRPL11-transduced progenitors (left) and quantifications ±SEM of eIF5A^H^/eIF5A and eIF5A/actin are presented (right n=6). Levels in control cells are set at “1”. (**E**) The differentiation of shCTRL- and shRPL11-transduced progenitors to a myeloid (CD11b^+^) versus erythroid (GlyA^+^) fate was evaluated by flow cytometry at day 3 of rEPO-induced differentiation. Representative CD11b/GlyA dot plots (left) and quantifications are presented (right, n=10). (**F**) OCRs of shCTRL- and shRPL11-transduced progenitors were evaluated at day 1 of differentiation and representative data (left, n=3-6 technical replicates) and mean basal OCR levels ± SEM are presented (right, n=3 independent experiments). *p<0.05; **p<0.01; ***p<0.001; ***p<0.0001; ns, not significant

We further assessed hypusination in CD34^+^ progenitors from del(5q) MDS patients. CD34^+^ progenitors were cytokine-expanded for 4 days and hypusination was then evaluated before and after rEPO-induced erythroid differentiation (day 0 and day 3). Similar to *Rps14*-haploinsufficient murine progenitors, the del(5q)-MDS progenitors exhibited high expression of eIF5A but hypusination levels were lower. Moreover, following rEPO-induced differentiation, there was a significant loss of eIF5A^H^ in del(5q)-MDS progenitors, accompanied by an ineffective erythroid differentiation (**Fig. 7C** and **S7B**). Thus, in both murine and human progenitors, the ineffective erythropoiesis linked to haploinsufficiency of RPS14 is associated with a loss of hypusinated eIF5A.

To specifically evaluate the link between RPs and hypusination as a function of erythroid differentiation, we downregulated expression of *RPL11*, a RP gene wherein mutations lead to a dramatic decrease in progenitor cell proliferation and a delayed erythroid differentiation (Flygare et al., 2005; Moniz et al., 2012). shRNA-mediated knockdown of *RPL11* resulted in a 90% decrease in expression and as previously reported, a significantly reduced progenitor cell proliferation (p<0.0001, **Fig. 7SC**). Strikingly, the shRNA-mediated downregulation of RPL11 resulted in a significant decrease in hypusination despite the maintenance of eIF5A levels (p<0.01, **Fig. 7D**). Moreover, the attenuated hypusination resulting from the downregulation of RPL11 was associated with a loss in rEPO-induced erythroid differentiation and almost 50% of progenitors underwent a CD11b myeloid differentiation fate (**Fig. 7E** and **S7D**). Finally, the impact of RPL11 expression on hypusination and erythroid commitment directly correlated with the mitochondrial metabolism of the cells. RPL11-downregulated progenitors, with an attenuated hypusination profile, demonstrated a significant loss in oxidative phosphorylation (**Fig. 7F** and **S7E**). Thus, these data reveal a novel role for RPs in controlling eIF5A hypusination in erythroid progenitors, resulting in the proper coordination of mitochondrial translation and metabolism and subsequent erythroid differentiation.

## DISCUSSION

Translation is a critical component in the regulation of multiple developmental pathways, cell activation, and cancer progression (Fabbri et al., 2021; Sanchez et al., 2016; Sonenberg and Hinnebusch, 2007; Tahmasebi et al., 2018). Within the broad realm of protein synthesis, hypusinated eIF5A while stimulating general translation, also drives specific translational programs impacting cancer, diabetes, and infectious diseases (Dever et al., 2014; Hoque et al., 2017; Kaiser, 2012; Tauc et al., 2021). Notably though the impact of eIF5A in erythropoiesis, a process that is particularly sensitive to translational regulation, has not been evaluated. Indeed, defects in protein translation selectively perturb the commitment of hematopoietic stem cells to the erythroid lineage. The data that we present here show that optimal protein synthesis in erythroid progenitors is dependent on SLC7A1-mediated arginine uptake, resulting in the hypusination of eIF5A by the arginine-catabolized spermidine moiety. Under conditions where hypusination is suboptimal, hematopoietic progenitors are not capable of undergoing erythroid differentiation and are skewed to a myeloid lineage fate. Thus, our study reveals a direct link between arginine metabolism and eIF5A-induced protein synthesis in controlling the erythroid-myeloid lineage commitment of human hematopoietic stem-progenitor cells. Furthermore, we uncover a critical role for eIF5A activity in the defective erythropoiesis associated with ribosomopathies.

Research uncovering the importance of hypusinated eIF5A in physiological as well as pathological processes has grown rapidly in the past few years. Following the purification of eIF5A from rabbit reticulocytes (Kemper et al., 1976), this factor has been identified as required for the translation elongation process rather than the initiation process (Saini et al., 2009), especially important for facilitating the translation of proteins containing polyproline residues and other motifs that are difficult for the ribosome to synthesize (Doerfel et al., 2013; Gutierrez et al., 2013; Ude et al., 2013). Notably, it is the only protein in eukaryotes and archaea to contain the amino acid hypusine [*N*^ε^-(4-amino-2-hydroxybutyl)lysine], a modification that is essential for its activity (Park and Wolff, 2018; Saini et al., 2009; Wolff et al., 2007). eIF5A is required for the growth of *S. cerevisiae* (Chattopadhyay et al., 2008) and has also been shown to be highly expressed at all mouse embryonic stages, with hypusination required for muscle differentiation (Parreiras-e-Silva et al., 2010; Tauc et al., 2021). Moreover, in humans, heterozygous variants in *EIF5A* have recently been shown to cause a syndrome characterized by developmental delay, microcephaly, and congenital malformations (Faundes et al., 2021). eIF5A is also important in regulating host-pathogen interactions from both sides; augmented hypusine synthesis increases the virulence of *Fusarium graminearum*, a devastating fungal pathogen of cereals (Martinez-Rocha et al., 2016) and the *Helicobacter pylori*/*Citrobacter rodentium*-induced synthesis of hypusine in macrophages is required for their optimal anti-microbial responses (Martinez-Rocha et al., 2016). Indeed, within immune cell lineages, hypusination has been found to control the differentiation and function of B cells, macrophages, NK cells, and T cells (Alsaleh et al., 2020; O’Brien et al., 2021; Puleston et al., 2021; Puleston et al., 2019; Zhang et al., 2019a).

The downstream effectors via which hypusinated eIF5A regulates cell differentiation and function are still being elucidated. While experimental studies in yeast as well as mammalian cells have identified an essential role for eIF5A in all translation (Kang and Hershey, 1994; Kemper et al., 1976; Mandal et al., 2016; Manjunath et al., 2019), hypusination is sensitive to the metabolic state of the cell and can also alter a cell’s metabolism. For reasons that are not completely clear, hypusinated eIF5A appears to have a skewed impact on mitochondrial protein translation, with defective hypusination resulting in decreased oxidative phosphorylation in macrophages and T cells (Puleston et al., 2021; Puleston et al., 2019) and increased reliance on glucose in kidney cells (Cougnon et al., 2021; Melis et al., 2017). Furthermore, the mitochondrial programming induced by hypusinated eIF5A, with increased levels of autophagy, can protect cell function––reversing B cell senescence (Zhang et al., 2019a), promoting vaccine immunogenicity in the elderly (Alsaleh et al., 2020), and protecting the brain from premature aging as well as ischemic-related stress (Bourourou et al., 2021; Liang et al., 2021b).

The findings that we present here demonstrate the critical nature of eIF5A-dependent mitochondrial function in regulating the erythroid commitment of HSPCs. Early erythroid differentiation requires mitochondrial function (Gonzalez-Menendez et al., 2021; Liu et al., 2017; Luo et al., 2017; Oburoglu et al., 2014; Zhao et al., 2016) and we now find that this metabolic state is dependent on eIF5A function. In the absence of eIF5A activity, mitochondrial complex I, II, III, IV and IV enzymes were decreased by greater than 80%, coupled to an almost complete loss of oxygen consumption. The impact of eIF5A on mitochondrial function directly regulated erythropoiesis as the abrogated erythroid commitment of DHPS-attenuated progenitors could be partially rescued by succinate; succinate significantly increased oxidative phosphorylation in eIFA-abrogated conditions, resulting in a 2-fold increase in the differentiation of GlyA^+^ erythroblasts. Thus, we demonstrate that eIF5A-dependent OXPHOS is a *sine qua non* for erythroid differentiation. Nonetheless, the mechanisms explaining how this cytoplasmic translation elongation factor impacts mitochondrial protein translation in diverse cell/organ systems remains to be determined.

The rapid initiation of translation allows erythroid progenitors to immediately respond to environmental cues such as heme and iron availability as well as oxygenation, stimulating erythroid differentiation programs (Moore and von Lindern, 2018; Vatikioti et al., 2019). It is therefore not surprising that multiple translation initiation factors such as eIF4G, eIF2α, EIF4EBP1 have been found to play important roles in erythropoiesis (Alvarez-Dominguez et al., 2017; Paolini et al., 2018; Tiu et al., 2021; Zhang et al., 2019b; Zhang et al., 2018). Moreover, mutations in different ribosomal proteins underlie bone marrow failure syndromes that are associated with anemia. Ribosomal protein haploinsufficiency is responsible for Diamond-Blackfan anemia, a disorder characterized by hypoplastic, macrocytic anemia, as well as for the macrocytic anemia exhibited by patients with 5q-deleted myelodysplastic syndrome (Chiabrando and Tolosano, 2010; Narla and Ebert, 2010). These patients exhibit general decreases in protein synthesis but more recent data indicate that shifts in ribosome availability impact the translation of specific transcripts that drive erythroid differentiation (Bello et al., 2018; Gazda et al., 2006; Horos et al., 2012; Khajuria et al., 2018; Mills and Green, 2017). In light of our data, it is therefore now opportune to assess the impact of ribosomal proteins on eIF5A-dependent translation, especially since numerous ribosomal proteins have been identified in a hypusine network interactome (Sievert et al., 2012). Our data reveal significantly attenuated hypusination of eIF5A in both murine and human hematopoietic progenitors with reduced expression of RPS14 and RPL11. Moreover, while translation of mitochondrial proteins has not been specifically evaluated in RP erythroblasts, our finding that reduced eIF5A activity in erythroblasts results in decreased mitochondrial function led us to evaluate mitochondrial activity in RP-modulated cells. Importantly, we find that a loss of eIF5A activity in RP-haploinsufficient or RP-downregulated progenitors is coupled to a defective mitochondrial metabolism. Thus, our study brings a novel perspective to our understanding of the defective protein synthesis characterizing ribosomopathies, extending defective translation to eIF5A-mediated processes.

The uncovered link between ribosomal proteins and eIF5A activity in erythroid differentiation opens new therapeutic perspectives for patients with ribosomopathies, including patients with Diamond-Blackfan anemia and del(5q)-MDS. L-leucine induces muscle protein synthesis (Norton and Layman, 2006), likely through the mTOR-mediated activation of the 5′cap-binding eIF4F complex (Drummond and Rasmussen, 2008; Sancak et al., 2008; Saxton and Sabatini, 2017; Son et al., 2020; Stipanuk, 2007). Notably, this amino acid has recently been found to increase protein synthesis in both DBA and del(5q) MDS erythroblasts, resulting in improved erythroid differentiation (Bello et al., 2018; Payne et al., 2012). Even more importantly, a recent phase I/II clinical study of leucine supplementation in DBA patients revealed an erythroid response in approximately 15% of patients and increased growth in almost half the patients (Vlachos et al., 2020). Interestingly though, the identity of the proteins impacted in these erythroblasts are not known. We hypothesize that the erythroid protein subsets regulated by the leucine-mediated induction of eIF4F complex (Saxton and Sabatini, 2017) may differ from those that are dependent on hypusinated eIF5A. eIF5A hypusination can potentially be increased by arginine supplementation, or more directly by spermidine itself. Indeed, arginine supplementation can improve endothelial dysfunction and immune responses, and possibly serve as an adjuvant to cancer treatments (Beal et al., 2019; Deveaux et al., 2016; Kim et al., 2018). In the context of erythroid differentiation, arginine was shown to increase hemoglobin levels in patients with renal disease (Tarumoto et al., 2007) but it was not efficacious in hydroxyurea-treated patients with sickle cell disease (Little et al., 2009), possibly because protein synthesis was not limiting. The polyamine spermidine may have greater impact than arginine in conditions where eIF5A activity is limiting as spermidine has been found to demonstrate therapeutic efficacy in several pathological conditions: *i)* protecting brain function and cognition in aging *Drosophila melanogaster* and mice (Hofer et al., 2021; Liang et al., 2021b; Schroeder et al., 2021); *ii)* reversing B cell senescence (Zhang et al., 2019a); *iii)* improving vaccinal responses in the elderly (Alsaleh et al., 2020); and *iv)* rescuing the micrognathia phenotype caused by decreased eIF5A expression in a zebrafish model (Faundes et al., 2021). While the potential to develop a therapeutic strategy to deliver high doses of spermidine to humans is not yet clear, it is very promising to note that a prospective cohort study found a significant correlation between high nutritional spermidine intake in humans and cognitive function (Schroeder et al., 2021). The data that we present here support the clinical evaluation of a combined nutrient supplementation in patients with ribosomopathies, with the goal of promoting eIF4F- and eIF5A-dependent protein synthesis through leucine and spermidine, respectively.

## METHODS

### CD34^+^ cell isolation and *ex vivo* differentiation assays

All studies involving human samples were conducted in accordance with the declaration of Helsinki. CD34^+^ cells were isolated from umbilical cord blood (UCB) within 24h of vaginal delivery of full-term infants after informed consent and approval by the “Committee for the Protection of Persons” (IRB) of the University Hospital of Montpellier. Bone marrow (BM) aspirates were obtained from deidentified donors at the Clinic Hematology Department of the University Hospital of Montpellier (CHU, Montpellier, France) and Moffitt Cancer Center & Research Institute (Tampa, FL, USA), after informed consent and approval from the respective IRBs. MDS and non MDS patients’ samples promoted at the CHU of Montpellier were obtained from the HemoDiag prospective cohort of patients with hemopathy (NCT02134574). Approximately 100 UCB and 6 BM samples were used in this study. Samples did not contain any identifiers including sex, race or ethnic origin. All samples were split such that they were used in all experimental groups. Thus, there was no need to use any specific criteria to allocate biological samples to experimental groups. For the vast majority of experiments, at least 3 biological samples were used.

CD34^+^ cells were selected using the anti-hCD34 Microbead Kit (Miltenyi Biotech) following the manufacturer’s instructions. For MDS samples, the anti-hCD34 Microbead UltraPure human kit (Miltenyi Biotech) was used. Selected cells (1 × 10^5^ cells/ml) were expanded in StemSpan H3000 media (StemCell Technologies Inc) supplemented with StemSpam CC100 (StemCell Technologies Inc) at 37°C with a humidified 5% CO2 atmosphere. Cells were plated in 24-well or 48-well *Nunclon Delta surface plates (Nunc, Thermo Fisher Scientific)*. After 4 days of expansion, cells were differentiated in IMDM medium (Invitrogen) supplemented with human holo-transferrin (200 mg/L, Sigma-Aldrich), human insulin (10 mg/L, Sigma-Aldrich), rhuSCF (10 ng/ml, Amgen), rhuIL-3 (1 ng/ml, R&D Systems). Erythropoiesis was induced by addition of 3 U/ml recombinant human erythropoietin (rEPO; Eprex, Janssen-Cilag). CD34^+^ cells were not expanded, and erythropoiesis was directly induced after selection in Supplementary Fig 2A. Cytokines were supplemented every 3-4 days, with media changes. Cells were centrifuged at 400g for 4min at RT and then resuspended in fresh media to maintain cell concentrations between 0.1 and 1.5 × 10^6^ cells/ml. For arginine deprivation experiments, IMDM for SILAC (Thermo Fisher Scientific) was used supplementing the corresponding concentration of lysine (0.8 mM, Sigma-Aldrich). Arginine (0.40mM, Sigma-Aldrich) was added in the control cells. In some experiments, progenitors were differentiated in the presence of 1 mM difluoromethylornithine (DFMO, provided by Dr. Thomas Dever), 5 µM N1-Guanyl-1,7-diaminoheptane (GC7, SantaCruz Biotechnology), 10 µM N1,N11-Diethylnorespermine (DENS, Tocris), 100 µM Trans-4-Methylcyclohexylamine (MCHA, Sigma-Aldrich), 100 µM N-(3-Aminopropyl)cyclohexylamine (APCHA, SantaCruz Biotechnology), 2.5 µM AMXT-1501 tetrahydrochloride (MedChemExpress), 5 µM ciclopirox (CPX, Sigma-Aldrich), 3 mM 2-Imino-1-imidazolidineacetic acid (cyclocreatine, Sigma-Aldrich), 50 µM *N*^*G*^,*N*^*G*^-Dimethylarginine dihydrochloride (ADMA, Sigma-Aldrich), 5 mM chloramphenicol (CPL, Sigma-Aldrich), 5 mM diethylsuccinate (Suc, Sigma-Aldrich). 100 µM of the polyamines putrescine or spermidine (Put or Spd, Sigma-Aldrich) were added in cells treated with DMFO alone or with DFMO and AMXT-1501. Drugs were freshly added after every media change. As chloramphenicol was dissolved in ethanol and ciclopirox in DMSO, control conditions included cultures using the same concentration of ethanol or DMSO (0.5% and 0.025%, respectively). Since cyclocreatine was dissolved in 1mM HCl, the pH of the medium was restored with 1mM NaOH.

### Murine progenitor assays

*Rps14*^+/*-*^*Mx1Cre*^+^ and *Mx1Cre* murine hematopoietic progenitor cell lines were cultured in RPMI-1640, supplemented with 10%FBS, GlutaMAX, Penicillin-Streptomycin, rmuSCF (50 ng/ml) and β-estradiol (0.5 mM). Cells were passaged every 3-4 days at a 1:5 dilution.

### Virus production and transduction of CD34^+^ progenitor cells

Lentiviral pLKO.1 plasmids harboring shRNAs directed against *SLC7A1* (ID: TRCN0000042967) and *DHPS* (ID: TRCN0000330717 and ID: TRCN0000330796) were obtained from Sigma-Aldrich. Cells were transduced with the shRNA plasmids where the 714bp sequence encoding for EGFP was inserted in place of the puromycin gene at the unique BamHI and KpnI restriction sites. shRNA against *RPL11* has been previously reported (Flygare et al., 2005; Moniz et al., 2012), sequences were synthesized (Eurogentec), and cloned into a pBlue Script containing the H1 promoter (Agilent). The casettes were inserted into a lentiviral vector pRRLsin-PGK-eGFP-WPRE (Addgene plasmid # 12252, a kind gift of D. Trono). Virions were generated by transient transfection of 293T cells (2-3×10^6^ cells/100 mm plate in 7 ml DMEM) with these vectors (10 µg) together with the Gag-Pol packaging construct PsPax2 (5 µg) and a plasmid encoding the VSV-G envelope, pCMV-VSV-G (2.5 µg), as described previously (Loisel-Meyer et al., 2012). Cells were transfected for at least 18 hours, and transfection efficiency was verified by monitoring GFP fluorescence by microscopy. Viral supernatant was harvested 24 hours post-transfection and virions were concentrated by overnight centrifugation at 4°C at 1,500g (with no break). Virions were resuspended in approximately 20 µl of RPMI with 10% FBS (per plate of cells), aliquoted and stored at -80°C. Titers were determined by serial dilutions of virus preparations on Jurkat cells and are expressed as Jurkat transducing units (TU/ml).

Prior to transduction of CD34^+^ hematopoietic stem and progenitor cells (HSPCs), cells were expanded for 3 days in StemSpan medium (Stem cell Technologies Inc) supplemented with 5% fetal bovine serum (FBS), 25 ng/ml rhuSCF (Amgen), 10 ng/ml rhuIL-3 and 10 ng/ml rhuIL-6 (R&D) at 37°C with a humidified 5% CO2 atmosphere. 5 × 10^5^ cells were then exposed to viral supernatants containing 4.8 × 10^5^ TU to 9.9 × 10^5^ TU (representing a multiplicity of infection of 1-2). After 72h of transduction, CD34^+^ cells (1 × 10^5^ cells/ml) were cultured in presence or absence of rEPO (3 U/ml). Transduction efficiency was monitored as a function of GFP expression at 72h post-transduction. For some experiments, GFP^+^ positive cells were sorted after-transduction.

### Flow cytometry

Expression of the CD34, CD36, CD49d, CD71, Glycophorin A, IL3R and CD11b surface markers was monitored on 1×10^5^ cells using the appropriate fluorochrome-conjugated monoclonal antibodies at a 1:100 dilution (mAb), except for CD36-APC and CD71-AF750 mAbs that were used at a 1:200 and 1:500 dilution, respectively (Beckman Coulter, Becton Dickinson, eBiosciences), in a total volume of 100 µl as previously described (Hu et al., 2013; Schulz et al., 2019). Cells were incubated in the dark for 20 minutes in PBS containing 2% FBS at 4ºC and then washed once in the same medium at 400g for 4 min prior to evaluation. Surface GLUT1 and SLC7A1 expressions were monitored by binding to their retroviral envelope ligand (RBD) fused to eGFP for GLUT1 (1:25 dilution in 50 µls; Metafora biosystems) or to the RBD-rFc fusion protein for SLC7A1 (1:25 dilution in 50 µl) for 30 minutes in PBS containing 2% FBS at 37ºC (Ivanova et al., 2017; Kim et al., 2004; Kinet et al., 2007; Manel et al., 2003; Swainson et al., 2005). RBD-rFc fusion protein incubation was followed by staining with an Alexa Fluor 488 or Alexa Fluor 647-coupled anti-rabbit IgG antibody (1:500 dilution in 100 µl, Thermo Fisher Scientific) for 20 minutes in PBS-2% FBS at 4ºC. Mitochondrial biomass was assessed in a total volume of 100 µl by MitoTrackerGreen or MitoTrackerDeepRed staining (20 nM; Invitrogen, Molecular Probes), while mitochondrial transmembrane potential levels were monitored by staining with MitoTrackerRed CMXRos (50 nM; Invitrogen, Molecular Probes). Incubations were performed in the dark for 20 minutes in PBS + 2% FBS at RT. Enucleation was evaluated by staining with the SYTO16 nucleic acid stain (1 μM, Invitrogen, Molecular Probes) for 15 minutes in PBS + 2% FBS at RT. Cell sorting was performed on a FACSARIA high-speed cell sorter (BD Biosciences) and analyses were performed on a FACS-Canto II (BD Biosciences) cytometers. A minimum of 10,000 events were recorded for each staining. Data analyses were performed using FlowJo software (BD Biosciences).

### Protein synthesis analyses

Protein synthesis rate was monitored by the Click-iT Plus OPP Protein Synthesis Assay (Molecular Probes, Thermo Fisher Scientific). 200,000 cells were incubated with 20 µM Click-iT OPP (O-propargyl-puromycin) in 100 µl of complete medium for 30 min at 37ºC and 5% CO2. Pre-treated cells with 1 µM cycloheximide (CHX, Sigma-Aldrich) for 24 hours were used as a control of translation inhibition. Cells were fixed with 2% paraformaldehyde for 10 min and permeabilized with 0.25% Triton X-100 for 5 min. Staining was performed following manufacturer’s protocol in 100 µl of reaction volume.

### Western blots

Cells were collected at the indicated time points and washed twice in PBS (400g for 4 min). Total protein content was extracted in RIPA lysis buffer (2.5 mM Tris-HCl pH 7.4, 1.3 mM NaCl, 0.01% NP-40, 0.5% sodium deoxycholate, 0.1% SDS, 1 mM EDTA) supplemented with protease inhibitors (10 mM NaF, 2 µg/ml aprotinin A) for 15 min at 4ºC (20 µl for 1×10^6^ cells). Samples were sonicated for 5 seconds at 40 kHz, and then centrifuged at 15,000g for 15 min at 4ºC. Supernatant was recovered and samples were prepared with 1:5 loading buffer (125mM Tris pH 6.8, 10% SDS, 20% Glycerol, 0.1% Bromophenol blue, 10% b-mercaptoethanol). Extracts were boiled for 5 min at 95ºC or heated for 2 minutes at 65ºC (OXPHOS complexes). Extracts corresponding to 5×10^5^ cells were loaded into a NuPAGE™ 4 to 12%, Bis-Tris, 1.0 mm, Mini Protein gel (Invitrogen, Thermo Fisher Scientific), and electrophoresis was run at 120V. Proteins were electrophoretically transferred to PVDF membranes (Thermo Fisher Scientific) for 2 hours at 250mA. Transfer efficiency was confirmed by Amido Black staining (Sigma-Aldrich). Membranes were incubated with 5% nonfat-dry milk in PBS-Tween 0.05% for 1 hour at RT. Blots were then incubated overnight at 4ºC with primary antibodies diluted in 3% nonfat-dry milk in PBS-Tween (1:2,000 Arginase 1; 1:2,000 Arginase 2; 1:2,500 anti-hypusine; 1:5,000 anti-eIF5A; 1:1,000 anti-OxPhos Rodent; 1:10,000 beta-actin), and for 1 hour at RT with a peroxidase anti-rabbit or anti-mouse immunoglobulin (1:5,000-1:20,000; 3% nonfat-dry milk in PBS-Tween). In some cases, blots were incubating in stripping buffer (1.5% Glycine, 0.1% SDS, pH 2.2) for 15 min at RT to remove previous antibodies. Antibodies were visualized using the Pierce™ ECL or the Supersignal™ west Femto maximum sensitivity substrate (Thermo Fisher Scientific), and images were registered using the Amersham Imager 680. Band intensities were quantified using ImageJ software (NIH). Ratios eIF5A^H^/eIF5A or protein/actin were represented. At least, 3 different biological samples were evaluated per condition.

### Quantitative Real Time PCR

Total RNA was isolated at the indicated time points and cDNA was synthetized using the RNeasy Mini Kit and the QuantiTect™ Reverse Transcription Kit (Qiagen) as per the manufacturer’s instructions. Quantitative PCR of cDNAs was performed using the Quantitect SYBR green PCR Master mix (Roche) with 10 ng of cDNA (by NanoDrop) and 0.5 µM primers in a final volume of 10 µl. Primer sequences are as follows: *SLC7A1*: 5’- CTATGGCGAGTTTGGTGC -3’ (forward) / 5’- CTATCAGCTCGTCGAAGGT -3’ (reverse); *DHPS*: 5’- GGGTTGGCCTTTGTATCTGA-3’ (forward) / 5’-TTTACAGGCCCAGATGAAGC-3’ (reverse); *RPL11:* 5’- GGGAACTTCGCATCCGCAA -3’ (forward) / 5’- CGCACCTTTAGACCCTTCTCC -3’ (reverse); and β-actin: 5’-GTCTTCCCCTCCATCGTG-3’ (forward) / 5’-TTCTCCATGTCGTCCCAG-3’ (reverse). Amplification of cDNAs was performed using the LightCycler 480 (Roche). Cycling conditions included a denaturation step for 5 min at 95°C, followed by 40 cycles of denaturation (95°C for 10 sec), annealing (63°C for 10 sec) and extension (72°C for 10 sec). After amplification, melting curve analysis was performed with denaturation at 95°C for 5 sec and continuous fluorescence measurement from 65°C to 97°C at 0.1°C/s. Each sample was amplified in triplicate. Data were analyzed by LightCycler® 480 Software (Version 1.5) and Microsoft Excel. Relative expression was calculated by normalization to β-actin as indicated (delta-Ct). Real-time PCR CT values were analyzed using the 2(Delta-Delta Ct) method to calculate fold expression (ddCt).

### Proteomics profiling

#### In-solution digestion

For proteomics analysis, 2×106 CD34^+^ cells were incubated with or without 5 µM GC7 in presence of EPO for 48 hours. Three different biological samples were evaluated for each condition. Cells were rapidly washed in ice-cold PBS, centrifuged twice (400g for 4 min), and dry pellets were conserved at -80ºC until extraction. Pellets were lysed for 30 minutes at 4°C in 8 M Urea, 75 mM NaCl, 50 mM Tris-HCl pH 8.0, 1 mM EDTA, 2 µg/ml aprotinin (Sigma), 10 µg/ml leupeptin (Roche), and 1 mM phenylmethylsulfonyl fluoride (PMSF) (Sigma). Protein concentration of cleared lysate was estimated with a bicinchoninic acid (BCA) assay (Pierce). Protein disulfide bonds were reduced with 5 mM dithiothreitol (DTT) at room temperature for 1 h, and free thiols were alkylated in the dark with 10 mM iodoacetamide (IAM) at room temperature for 45 min. The urea concentration in all samples was reduced to 2 M by addition of 50 mM Tris-HCl, pH 8.0. Denatured proteins were then enzymatically digested into peptides upon incubation first with endoproteinase LysC (Wako Laboratories) at 25 °C shaking for 2 h and then with sequencing-grade trypsin (Promega) at 25 °C shaking overnight, both added at a 1:50 enzyme:substrate ratio. Digestion was quenched via acidification to 1% formic acid (FA). Precipitated urea and undigested proteins were cleared via centrifugation, and samples were desalted using 50 mg tC18 1cc SepPak desalt cartridges. Cartridges were conditioned with 100% Acetonitrile (MeCN), 50% MeCN/0.1% FA, and 0.1% trifluoroacetic acid (TFA). Samples were loaded onto the cartridges and desalted with 0.1% TFA and 1% FA, and were then eluted with 50% MeCN/0.1% FA. Eluted samples were frozen and dried via vacuum centrifugation.

#### TMT labeling of peptides

Desalted peptides were reconstituted in 30% MeCN/0.1% FA and the peptide concentration was quantified with a BCA assay. With 50 µg peptide input per channel, samples were labeled with a TMT11 isobaric mass tagging reagent (Thermo) as previously described (Zecha et al., 2019). Samples were reconstituted in 50 mM HEPES, pH 8.5, at a peptide concentration of 5 mg/ml. Dried TMT reagent was reconstituted in 100% anhydrous MeCN at a concentration of 20 µg/µl, added to each sample at a 1:1 TMT:peptide ratio, and allowed to rea for 1 h at 25 °C. Labeling was quenched upon addition of 5% hydroxylamine to a final concentration of 0.25%, incubating for 15 min at 25 °C. TMT-labeled samples were combined, frozen, and dried via vacuum centrifugation. This dried sample was reconstituted in 0.1% FA and desalted using a 50 mg tC18 1cc SepPak cartridge as described above. The eluted sample was frozen and dried via vacuum centrifugation.

#### Basic Reverse Phase (bRP)

*Fractionation:* Labeled and combined peptides for proteome analysis were fractionated using offline basic reverse-phase (bRP) fractionation as previously described (Mertins et al., 2018). The sample was reconstituted in 900 µl bRP solvent A (2% vol/vol MeCN, 5 mM ammonium formate, pH 10.0) and loaded at a flow rate of 1 ml/min onto a custom Zorbax 300 Extend C18 column (4.6 × 250 mm, 3.5 µm, Agilent) on an Agilent 1100 high pressure liquid chromatography (HPLC) system. Chromatographic separation proceeded at a flow rate of 1 ml/min with a 96 min gradient, starting with an increase to 16% bRP solvent B (90% vol/vol MeCN, 5 mM ammonium formate, pH 10.0), followed by a linear 60 min gradient to 40% that ramped up to 44% and concluded at 60% bRP solvent B. Fractions were collected in a Whatman 2 ml 96-well plate (GE Healthcare) using a horizontal snaking pattern and were concatenated into 24 final fractions for proteomic analysis. Fractions were frozen and dried via vacuum centrifugation.

#### Liquid Chromatography and Mass Spectrometry

After reconstitution of dried fractions in 3% MeCN/0.1% FA to a concentration of 1 μg/μl, samples were analyzed via coupled Nanoflow LC-MS/MS using a Proxeon Easy-nLC 1000 (Thermo Fisher Scientific) and a Q-Exactive Plus Series Mass Spectrometer (Thermo Fisher Scientific). One μg of each fraction was separated on a capillary column (36-μm outer diameter × 75-μm inner diameter) containing an integrated emitter tip and heated to 50°C, followed by packing to a length of approximately 30 cm with ReproSil-Pur C18-AQ 1.9 μm beads (Maisch GmbH). Chromatography gradient consisted of solvent A (3% MeCN/0.1% FA) and solvent B (90% MeCN/0.1% FA), and the profile was 0:2, 1:6, 85:30, 94:60, 95:90, 100:90, 101:50, and 110:50 (minutes/percentage solvent B). The first 6 steps were performed at a flow rate of 200 nl/min and the last 2 at 500 nl/min. Ion acquisition was performed in data-dependent mode, acquiring HCD-MS/MS scans at a resolution of 35,000 on the top-12 most abundant precursor ions in each full MS scan (70,000 resolution). The automatic gain control (AGC) target was set to 3×106 ions for MS1 and 5×104 for MS2, and the maximum inject time to 120 ms for MS2. The collision energy was set to 30, peptide matching was set to “preferred,” isotope exclusion was enabled, and dynamic exclusion time was set to 20 seconds.

#### Data Analysis

Mass spectrometry data was processed using Spectrum Mill (Rev BI.07.04.210, proteomics.broadinstitute.org). Extraction of raw files retained spectra within a precursor mass range of 750-6000 Da and a minimum MS1 signal-to-noise ratio of 25. MS1 spectra within a retention time range of +/- 60 s, or within a precursor m/z tolerance of +/- 1.4 m/z were merged. MS/MS searching was performed against a human UniProt database. Digestion parameters were set to “trypsin allow P” with an allowance of 4 missed cleavages. The MS/MS search included fixed modifications, carbamidomethylation on cysteine and TMT on the N-terminus and internal lysine, and variable modifications, acetylation of the protein N-terminus, oxidation of methionine, and TMT-hypusination of lysine. Restrictions for matching included a minimum matched peak intensity of 30% and a precursor and product mass tolerance of +/- 20 ppm. Peptide matches were validated using a maximum false discovery rate (FDR) threshold of 1.2%, limiting the precursor charge range to 2 to 6. Protein matches were additionally validated, requiring a minimum protein score of 0. Validated data was summarized into a protein-centric table and filtered for fully quantified hits represented by 2 or more unique peptides. Non-human contaminants and human keratins were removed.

#### Statistical approach

Each protein ID was associated with a log2-transformed expression ratio for every sample condition over the median of all sample conditions. After median normalization, a 2-sample moderated T test was performed on the data to compare treatment groups using an internal R-Shiny package based in the limma library. P-values associated with every protein were adjusted using the Benjamini-Hochberg FDR approach (Benjamini and Hochberg, 1995).

Gene Set Enrichment Analysis (GSEA) was performed using the WebGestalt (WEB-based Gene SeT AnaLysis Toolkit) 2019 web tool (Liao et al., 2019). Only the proteins significantly downregulated by GC7 were analyzed, using the log_2_-transformed ratios of GC7/control as input. Reactome was chosen as functional database applying the affinity propagation. Cytoscape software was employed for proteomics protein-protein interaction network analysis (Shannon et al., 2003), applying the STRING plugin (Szklarczyk et al., 2015) and MCODE (Bader and Hogue, 2003) for the clustering analysis.

### Seahorse analysis

Oxygen consumption rates (OCR) and extracellular acidification rates (ECAR) were measured using the Xfe-96 Extracellular Flux Analyzer (Seahorse Bioscience, Agilent). The calibration plate was filled with 200 µl Milli-Q water per well and kept at 37ºC overnight. At least 2 hours prior to the assay, water was replaced by 200 µl XF calibrant (Agilent)/ well. Culture plates were treated with 20 µl/well of 0.1mg/ml of Poly-D-Lysine (Sigma-Aldrich) for at least 1 hour at RT. Poly-D-Lysine solution was then removed and plates were washed twice with 200 µl Milli-Q water/well, dried, and kept at 4ºC until the assay was performed. For Mito Stress Test, cells (1 × 10^5^ - 2 × 10^5^) were placed in 50 µl XF medium (non-buffered DMEM containing 10 mM glucose and 2 mM GlutaMax, pH 7.3-7.4, Agilent), centrifuged in a carrier tray (300g for 3 min with no break), and incubated for 30 min at 37ºC. An additional 130 µl XF medium was added during the calibration time (20-30 min at 37ºC). Cells were monitored in basal conditions and in response to oligomycin (1 µM; Sigma-Aldrich), FCCP (1 µM; Sigma-Aldrich), rotenone (100 nM, Sigma-Aldrich) and antimycin A (1 µM; Sigma-Aldrich). The same procedure was followed for Glycolysis Stress Test in non-buffered DMEM without glucose and glutamine (Sigma-Aldrich). Cells were monitored in basal conditions and in response to D-glucose (10 mM, Sigma-Aldrich), oligomycin (1 µM, Sigma-Aldrich) and 2-deoxyglucose (2DG, 100 mM, Sigma-Aldrich). Wave software (Agilent) was employed for running the Seahorse assay and analysis. At least 3 independent experiments were performed with >3 technical replicates per experiment.

### Arginine and glutamine uptake assays

Cells (5 × 10^5^) were washed twice in 1.5 ml of PBS containing 2%FBS at 400g for 4 min at RT and starved for arginine or glutamine by incubation at 37°C in 30 µl arginine or glutamine-free IMDM for 30 min (Thermo Fisher Scientific and Sigma-Aldrich, respectively). Radiolabeled L-[3,4-3H]-Arginine monohydrochloride or glutamine-L-[3,4-^3^H(N)] (Perkin Elmer) was added to a final concentration of 8 and 2 µM respectively (2 and 0.5 μCi in a total volume of 50 µl). Cells were incubated for 10 min at room temperature, washed twice in cold PBS containing 2%FBS (1,000g, 3 min), and solubilized in 500 μL of a 0.1% SDS solution. Cells were incubated with 200 µM N-ethylmaleimide (NEM, Sigma-Aldrich) during the uptake to inhibit the SLC7A1-dependent arginine uptake. Radioactivity was measured in 4.5 ml liquid scintillation (Perkin Elmer) using a Hidex 300 SL counter. Each independent experiment was performed in triplicate.

### Statistical analyses

Data are represented as individual values or means. Error bars represent the standard error of the mean (SEM). Data were analyzed with Prism (GraphPad software). All groups followed normal distribution, and *P*-values were determined by one-way ANOVA (Tukey’s post-hoc test) or t-tests, as indicated in the corresponding text and figure legends. Two-tailed t-tests were used in all figures. Tests were paired or unpaired as indicated. All statistical details of experiments can be found in the figure legends. For proteomic analysis, data were median normalized and subjected to a 1-sample moderated t test using an internal R-Shiny package based in the limma R library. Benjamini-Hochberg FDR methods was used to correct for multiple testing. Significance is indicated as * p<0.05, **p<0.01, ***p<0.001 and ****p<0.0001 and ns, for non-significant.

## Supplemental Figures

**Figure S1.**
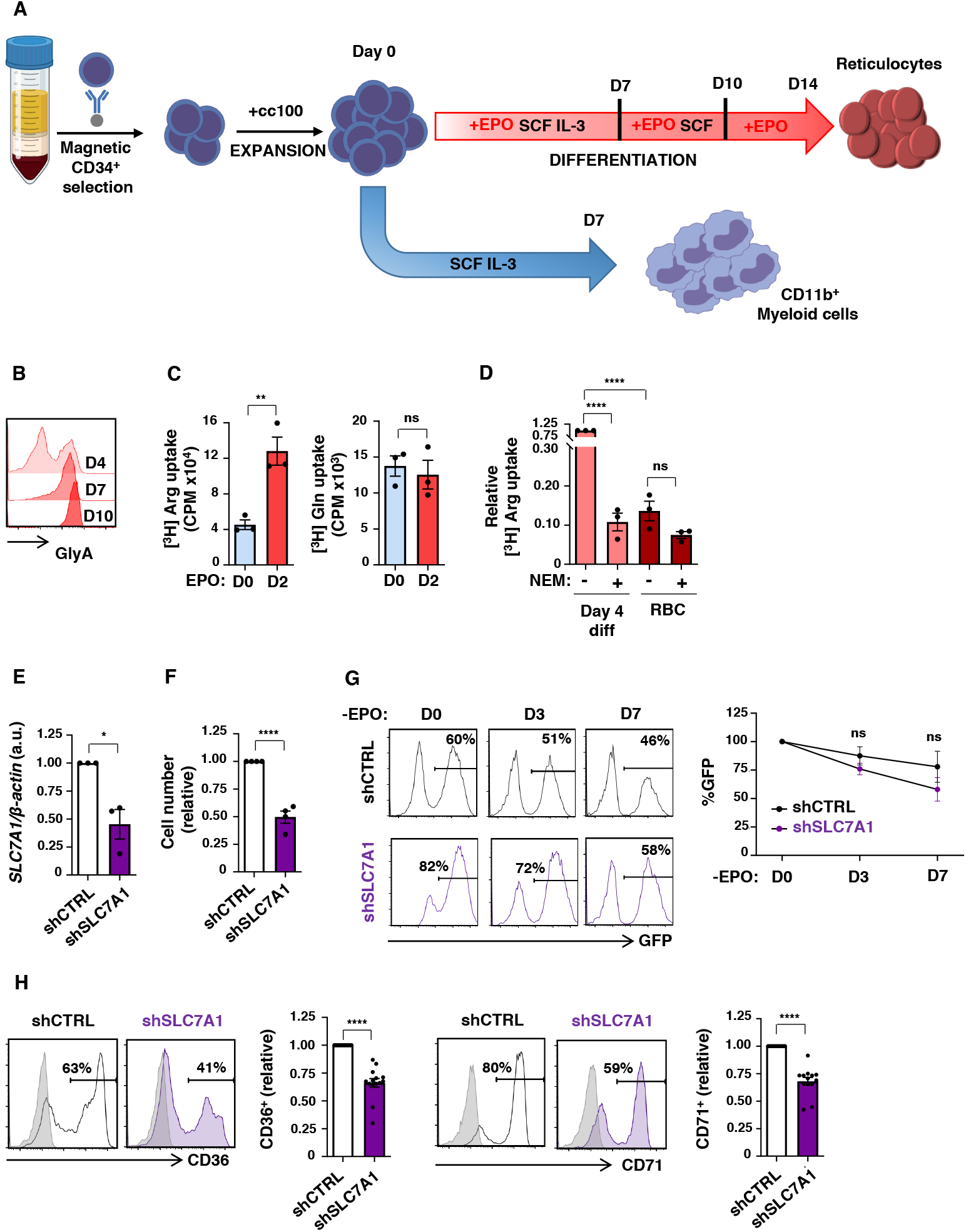
Selective disadvantage of SLC7A1/CAT1-downregulated progenitors upon erythroid but not myeloid differentiation. (**A**) Schematic representation of the *ex vivo* differentiation protocol of CD34^+^ progenitors in the presence of SCF/IL3/EPO or SCF/IL-3, resulting in erythroid and myeloid differentiation, respectively. (**B**) Expression of GlyA was evaluated at days 4, 7 and 10 of differentiation and representative histograms are shown. (**C**) Arginine and glutamine uptakes were monitored at days 0 and 2 of EPO-induced differentiation using [^3^H] L-arginine (2 µCi) or L-[3,4-^3^H (N)]glutamine (0.5 µCi), respectively, for 10 min at RT. Uptake is expressed as mean counts per minute (CPM) of triplicate samples ±SEM. (**D**) [^3^H] l-arginine uptake was monitored at day 4 of differentiation as well as in mature RBC, in the absence or presence of the N-ethylmaleimide (NEM) SLC7A1 inhibitor. Mean CPM of triplicate samples in 3 independent experiments ± SEM are presented. (**E**) Expression of *SLC7A1* was evaluated by qRT-PCR in shCTRL and shSLC7A1-transduced progenitors, sorted on the basis of GFP expression, and normalized to actin. Means ± SEM of 7 independent experiments are shown with values in control cells set at “1”. (**F**) Relative cells numbers were evaluated at day 3 of differentiation following transduction with an shCTRL or shSLC7A1 vector (n=4). (**G**) The evolution of shCTRL- and shSLC7A1-transduced progenitors was evaluated at days 0, 3, and 7 in the absence of EPO stimulation, as a function of GFP expression, and representative histograms are presented (left). Quantification of the loss of GFP expression is presented relative to day 0, set at “100%” (n=4, right). (**H**) CD36 and CD71 levels were evaluated in shCTRL- and shSLC7A1-transduced progenitors at day 3 of EPO stimulation. Representative histograms are shown and quantification of positive cells were compared relative to shCTRL cells (n=14 for CD36 and n=13 for CD71). *p<0.05; **p<0.01; ****p<0.0001; ns, not significant

**Figure S2.**
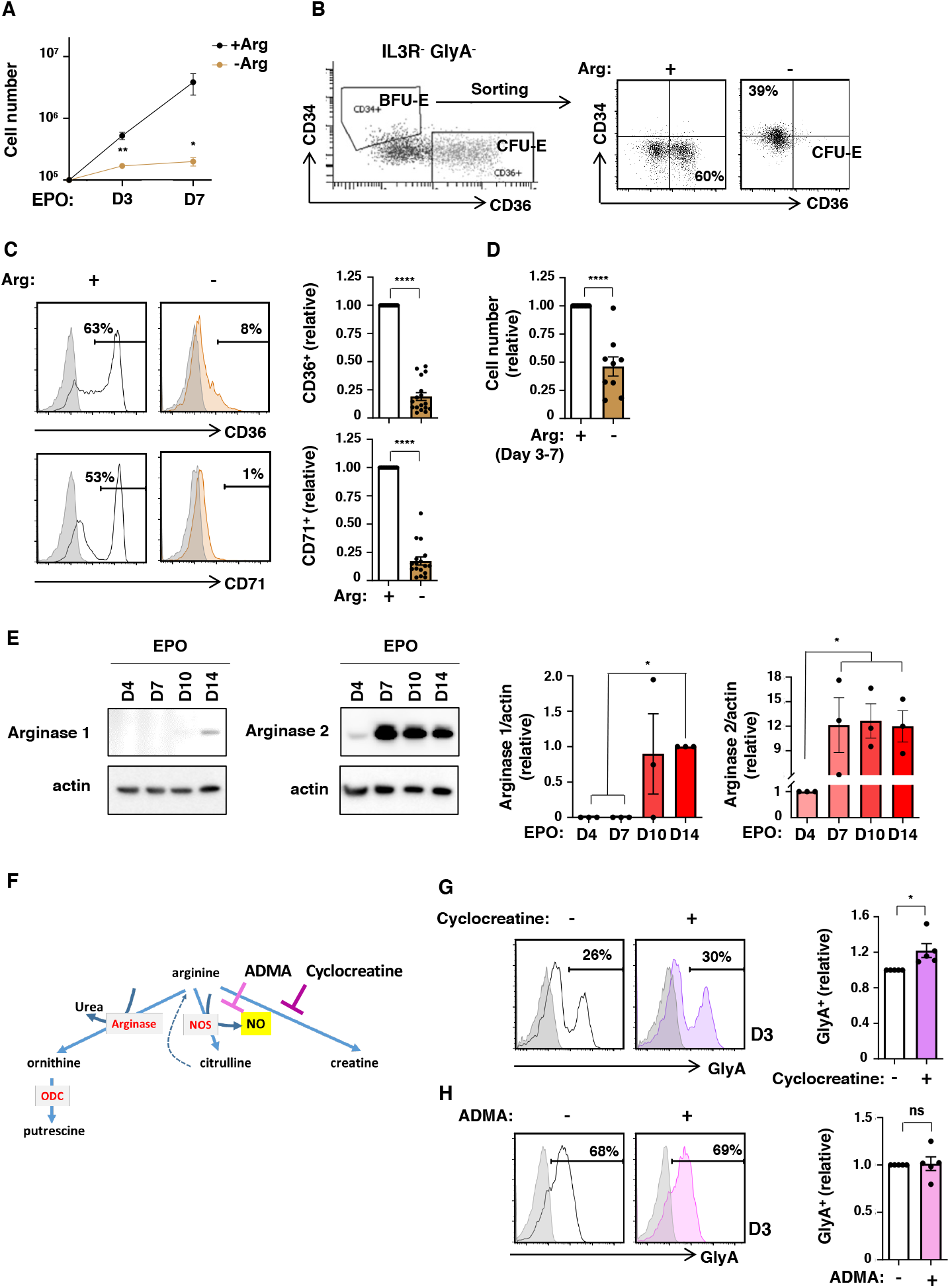
Erythroid lineage specification is not dependent on arginine catabolism to NO or creatine. (**A**) Proliferation of progenitors differentiated in EPO in the presence or absence of Arg is presented as mean cell numbers ± SEM at the indicated time points (n=8). (**B**) Representative dot plot showing CD34/CD36 profiles of IL3R^-^GlyA^-^ progenitors, distinguishing between CD34^+^CD36^+^ BFU-E and CD34^-^CD36^+^ CFU-E (left). Sorted BFU-E were differentiated in the presence or absence of arginine (Arg) and CD34/CD36 profiles were evaluated 4 days later (right). (**C**) CD36 and CD71 expression were evaluated at day 3 of EPO-induced differentiation in the presence or absence of Arg. Representative histograms (left) and quantification of relative levels (right, n=17) are presented. (**D**) Progenitors were differentiated for 3 days in the presence of EPO and differentiation was then continued in the presence or absence of Arg until day 7. Cell numbers relative to the presence or Arg are presented (n=9). (**E**) Expression of arginase 1 and 2 were evaluated at days 4, 7, 10, and 14 of erythroid differentiation and representative immunoblots as well as actin immunoblots are shown (left). Quantification of arginase expression relative to actin was evaluated, and levels of Arg1 at day 14 were arbitrarily set at “1”, and Arg2 levels at day 4 were arbitrarily set at “1” (n=3). (**F**) Schematic showing the generation of putrescine, citrulline, NO, and creatine from arginine. Enzymes are shown in red and cyclocreatine, an analog of the creatine and inhibitor of the creatine biosynthesis as well as asymmetric dimethylarginine (ADMA), a competitive inhibitor of nitric oxide synthase (NOS), were used to inhibit these respective pathways. (**G**) The impact of cylocreatine (top, 3mM) and ADMA (bottom, 50µM) on erythroid differentiation was evaluated as a function of GlyA staining at day 3 and representative histograms (left) as well as quantification of GlyA expression (n=5) are presented (right).*p<0.05; **p<0.01 ****p<0.0001; ns, not significant

**Figure S3.**
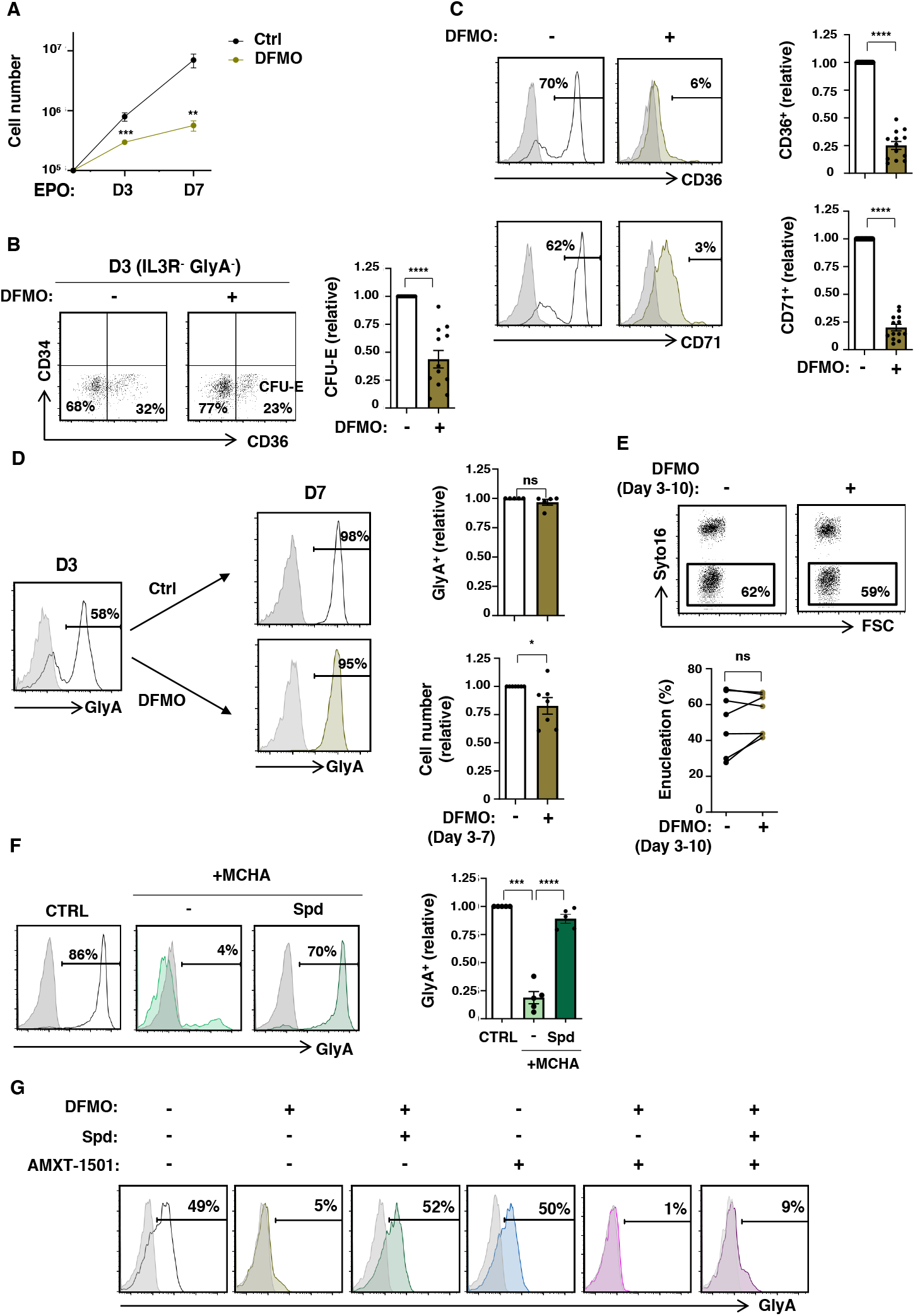
Spermidine rescues erythroid differentiation in the absence of polyamine biosynthesis. (**A**) Proliferation of progenitors differentiated in EPO in the absence or presence of 1mM DFMO is presented as mean cell numbers ± SEM at the indicated time points (n=13). (**B**) Representative dot plots of CD34/CD36 profiles of IL3R^-^GlyA^-^ cells are shown for progenitors differentiated in the absence or presence of DFMO (day 3) and the percentages of IL3R^-^GlyA^-^CD34^-^CD36^+^ (CFU-E) are indicated (left). Quantification of CFU-E relative to the absence of DFMO are presented (n=12, right). (**C**) Differentiation was monitored as a function of CD36 and CD71 expression in the presence or absence of DFMO and representative histograms are presented (left). Quantification of expression relative to control conditions are shown (right, n=13). (**D**) Progenitors were differentiated in the presence of EPO until day 3 and then differentiation was continued in the absence or presence of DFMO. Representative histograms showing GlyA expression at days 3 and 7 are presented (left). Quantification of GlyA expression (top right) and cell counts (bottom right) are presented. (**E**) Following EPO-induced differentiation between days 3 and 10, in the absence or presence of DFMO, enucleation was monitored as a function of Syto16 staining. Representative dot plots (top) and comparisons of enucleation levels in 7 independent experiments (bottom) are shown. (**F**) MCHA-treated progenitors were differentiated with rEPO in the presence of absence of Spermidine. Representative histograms (left) and quantifications (n=5, right) are presented. (**G**) Erythroid differentiation was induced in the presence or absence of DFMO, spermidine (100µM) and AMXT-1501 (2.5µM) and representative histograms showing GlyA expression at day 3 are presented. *p<0.05; **p<0.01; ***p<0.001; ****p<0.0001; ns, not significant

**Figure S4.**
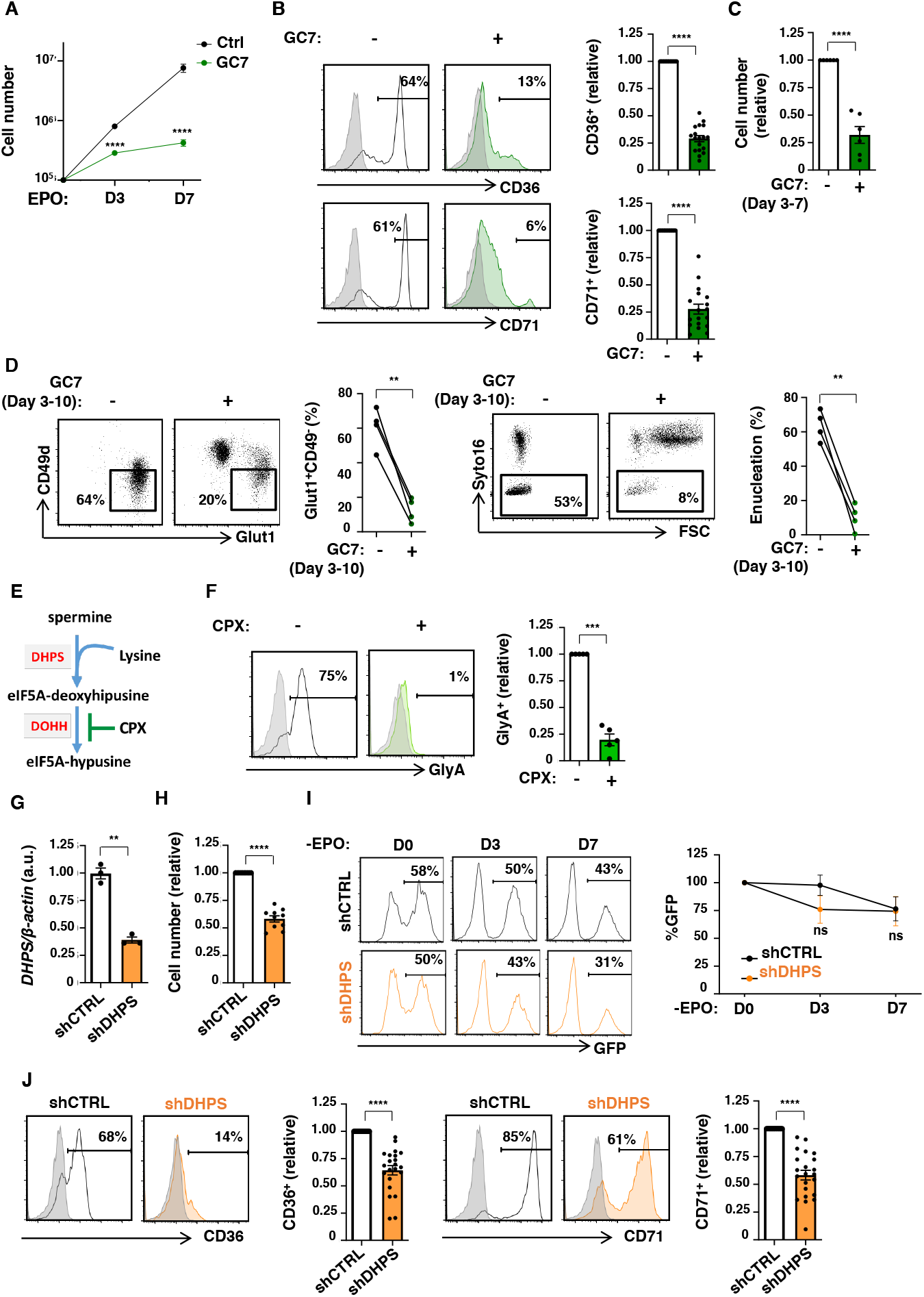
Myeloid differentiation is not inhibited by DHPS downregulation. (**A**) Proliferation of progenitors differentiated in EPO in the absence or presence of GC7 (5µM) is presented as mean cell numbers ± SEM at the indicated time points (n=17). (**B**) Erythroid differentiation was monitored as a function of CD36 and CD71 expression and representative histograms are shown (left). Quantification of expression relative to control conditions is presented (n=18, right). **(C)** Progenitors were differentiated for 3 days and then exposed to GC7 for days 3-7. Cell numbers at day 7, relative to control conditions are shown (n=6). (**D**) Differentiation of GlyA^+^ cells was monitored as a function of a CD49d^-^Glut1^+^ phenotype at day 10 and representative dot plots (left) as well as percentages in the absence or presence of GC7 are shown (left, n=8). Enucleation was monitored as a function of Syto16 staining (right, n=4). (**E**) Schematic showing the generation of eIF5a^H^ from spermine, as a function of the catalytic activities of DHPS and DOHH. Inhibition of DOHH by ciclopirox (CPX) is presented. (**F**) Erythroid differentiation in the absence or presence of CPX (5µM) was evaluated at day 3 of differentiation as a function of GlyA expression and representative histograms are shown (left). GlyA expression relative to control conditions is presented (n=5, right). (**G**) Expression of DHPS was evaluated by qRT-PCR in shCTRL- and shDHPS-transduced progenitors as a function of actin and means ± SEM of 3 independent experiments are shown. (**H**) Cell numbers were monitored at day 3 of differentiation and are presented relative to shCTRL-transduced progenitors (n=10). (**I**) Evolution of shCTRL- and shDHPS-transduced progenitors was monitored as a function of co-expressed GFP under conditions of non-erythroid (myeloid) differentiation. Representative histograms at the indicated time points are shown (left). Quantification of GFP expression relative to day 0 is shown (right, n=3). (**J**) CD36 (left) and CD71 (right) expression was monitored in shCTRL- and shDHPS-transduced progenitors at day 3 of differentiation and representative histograms are shown. Expression relative to controls conditions was evaluated and means ± SEM are presented (n=14). **p<0.01; ***p<0.001; ****p<0.0001

**Figure S5.**
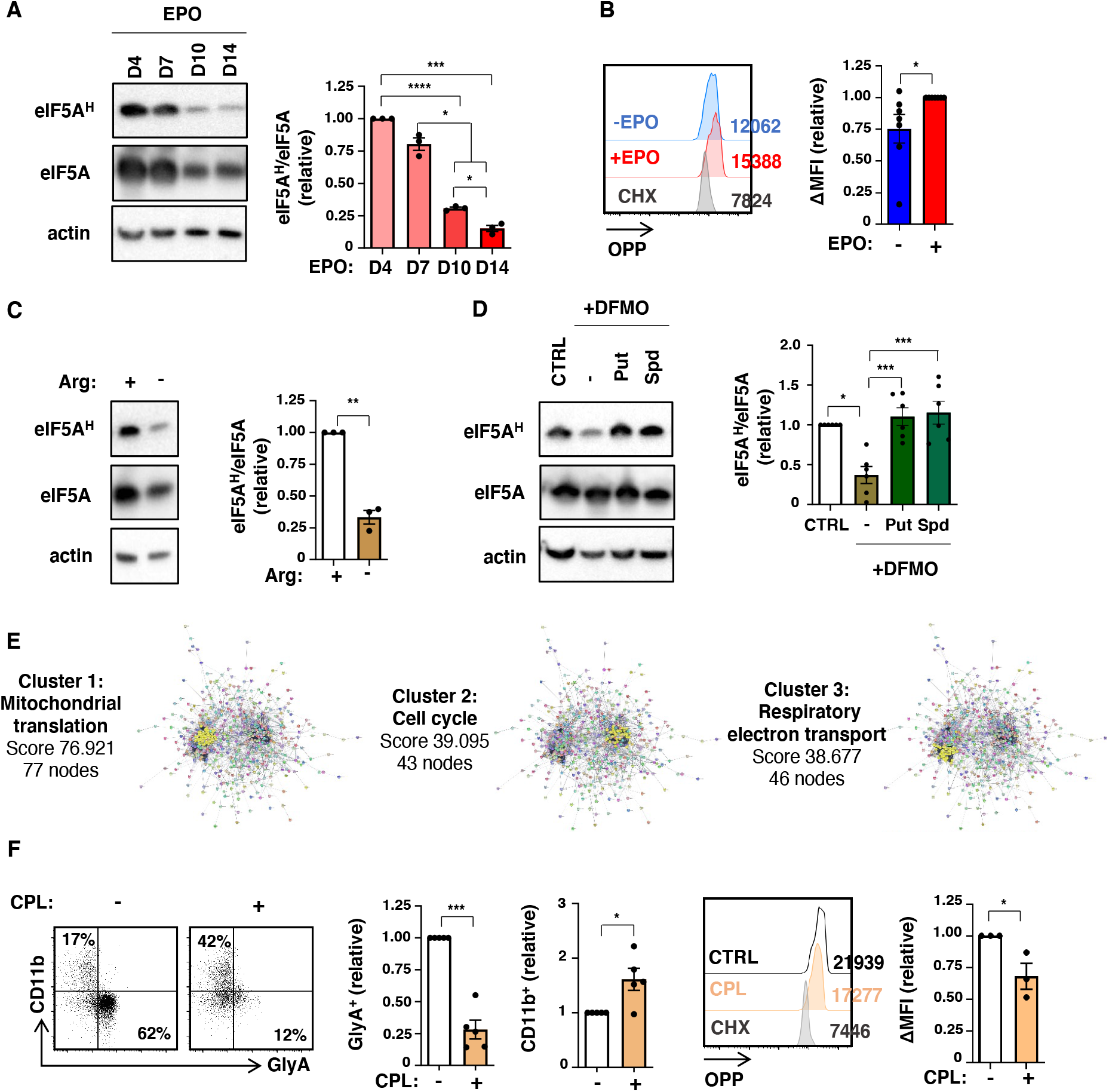
Arginine metabolism regulates hypusination of eIF5A in erythroid progenitors. (**A**) Hypusination of eIF5A was evaluated at days 4, 7, 10 and 14 of erythroid differentiation by immunoblotting with a polyclonal antibody against hypusinated human eIF5A (eIF5A^H^). Control immunoblots assessing total eIF5A protein and actin are presented (left). Quantification of hypusine relative to eIF5A was determined, with levels at day 4 arbitrarily set at 1 (n=3, right). (**B**) Protein synthesis was monitored by OPP and representative histograms in the presence or absence of EPO as well as in the presence of cycloheximide (CHX, 1µM) are presented and MFIs are indicated. Quantification of changes in OPP relative to EPO conditions are shown (n=7). (**C**) Hypusine levels were evaluated following differentiation in the presence or absence of arginine and immunoblots (left) as well as quantification (n=3, right) are shown. (**D**) Hypusination was evaluated at day 3 of erythroid differentiation in control conditions as well as in the presence of 1mM DFMO, either alone or supplemented with 100µM putrescine (Put) or spermidine (Spd). Representative immunoblots (left) and quantifications of eIF5A^H^ /eIF5A are shown (right, n=6). (**E**) PPI network clustering analysis identified mitochondrial translation, cell cycle and respiration electron transport pathways as significantly downregulated in the presence of 5µM GC7. Scores and nodes are indicated. (**F**) Erythroid differentiation was monitored at day 3 in the absence or presence of chloramphenicol (CPL, 5mM) as a function of CD11b/GlyA expression and representative dot plots and quantification (n=5) are presented (left). Protein expression in these conditions was monitored by OPP staining at day 1 of differentiation and representative histograms and quantification (n=3) are shown (right). *p<0.05; **p<0.01; ***p<0.001; ****p<0.0001; ns, not significative

**Figure S6.**
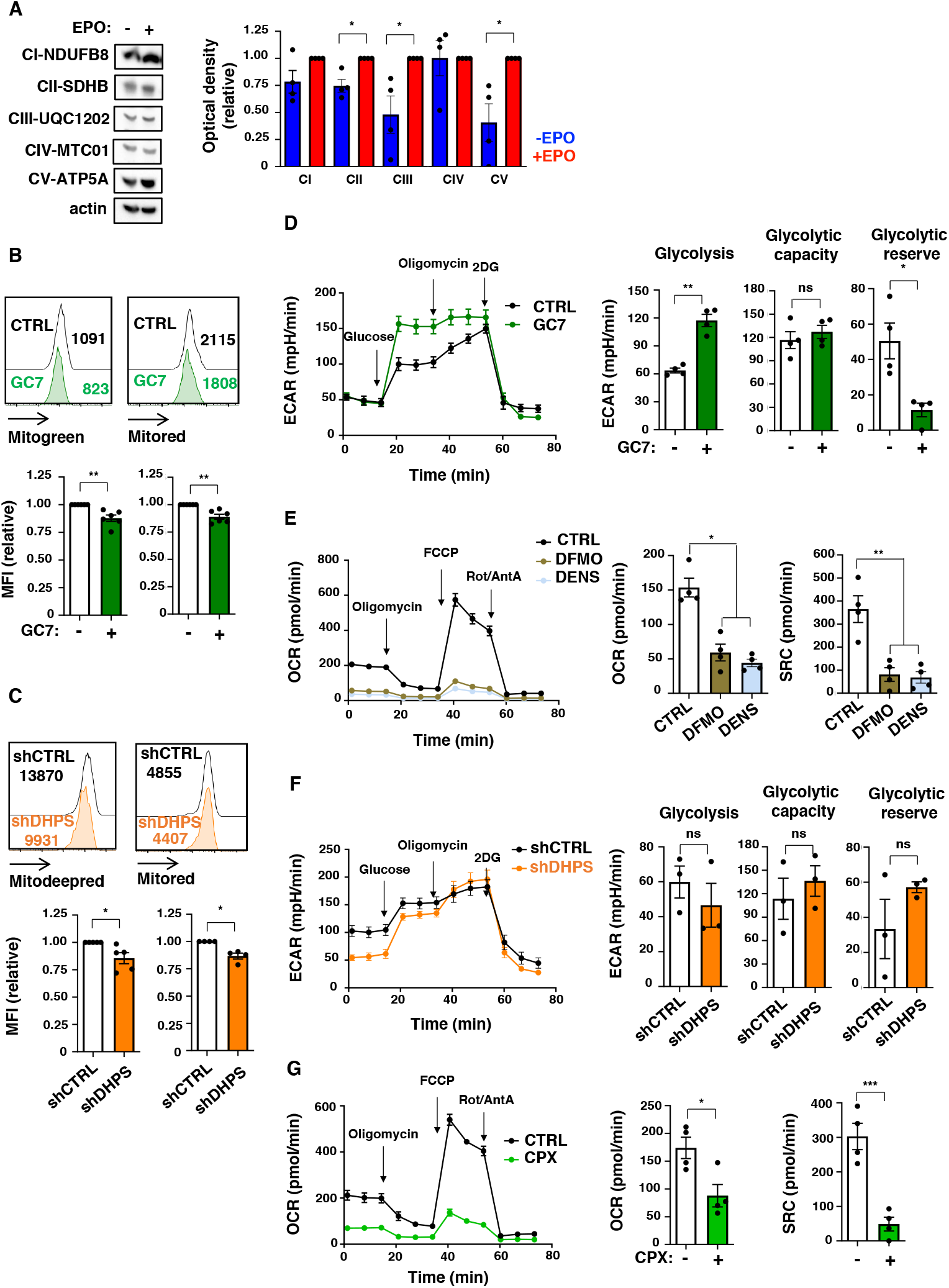
Hypusinated eIF5A regulates oxidative phosphorylation in hematopoietic progenitors. (**A**) Mitochondrial complexes (CI to CV) were monitored on progenitors differentiated for 4 days in the presence or absence of EPO. Representative immunoblots (left) and quantifications (relative to EPO-treated progenitors) are presented (n=4, right). (**B**) Mitochondrial biomass and polarization were monitored as a function of Mitotracker green and Mitotracker red staining, respectively, and representative histograms (top) and quantifications (bottom, n=6) are shown for progenitors differentiated in the absence or presence of GC7 (day 1). (**C**) shCTRL- and shDHPHS-transduced progenitors were stained with Mitotracker deepred and Mitrotracker red at day 1 of differentiation and representative histograms (top) as well as quantification of staining relative to shCTRL cells (bottom, n=4-5) are shown. (**D**) A glycolysis stress test was performed to evaluate the impact of 5µM GC7 treatment on glycolysis and a representative tracing at day 1 of differentiation is shown (left). Glycolysis, glycolytic capacity, and glycolytic reserve were evaluated using the Agilent Seahorse XF assay and means ± SEM are presented (means of 4 independent experiments performed with 3-6 technical replicates) (right). (**E**) OCR was monitored on day 1 of erythroid differentiation in the absence or presence of 1mM DFMO or 10µM DENS and a representative tracing (left) as well as means ± SEM of basal OCR and SRC in 4 independent experiments (with 3-6 technical replicates each) are presented (right). (**F**) A glycolysis stress test was performed on shCTRL- and shDHPS-transduced progenitors at day 1 of differentiation as in panel D and glycolysis, glycolytic capacity and glycolytic reserve are presented (n=3). A representative tracing (left) and means ± SEM are presented (right). (**G**) OCR was monitored on day 1 of erythroid differentiation in the absence or presence of 5µM CPX and means ± SEM of basal OCR and SRC in 4 independent experiments (with 3-6 technical replicates each) are presented (right). *p<0.05; **p<0.01; ***p<0.001; ns, not significant

**Figure S7.**
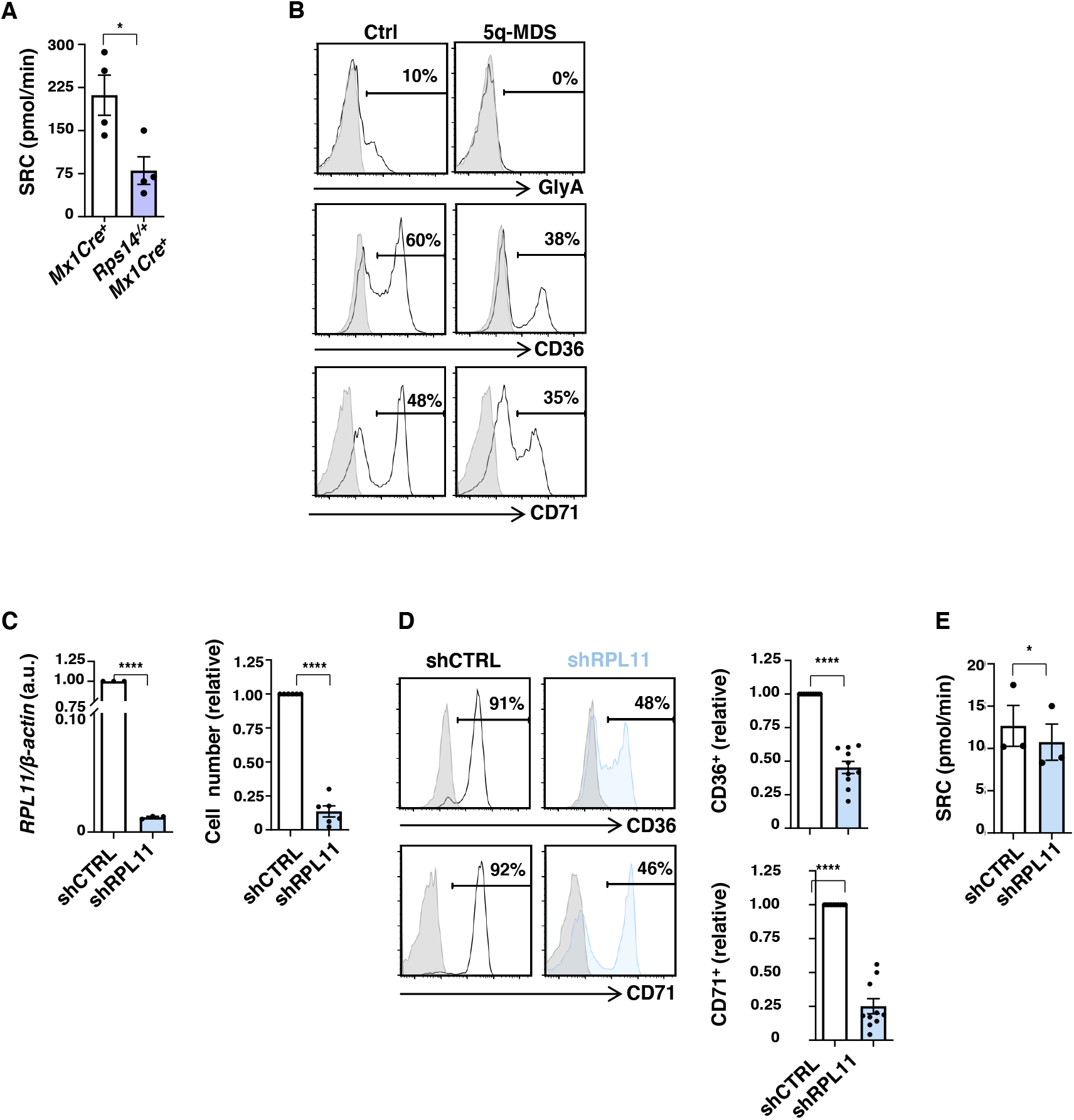
The defective erythroid differentiation associated with ribosomal protein insufficiency is linked to impaired hypusination. (**A**) SRC levels ± SEM (n=4) are presented for data presented in panel 7B in *Mx1Cre*^*+*^ and *Rps14*^*+/-*^*Mx1Cre*^*+*^ progenitors. (**B**) Expression of GlyA, CD36, and CD71 were evaluated in BM progenitors from a healthy control individual and a del(5q)-myelodysplastic syndrome patient at day 3 of erythroid differentiation. (**C**) Expression levels of RPL11 were evaluated by qRT-PCR following transduction with shRPL11 virus (left). mRNA levels were normalized to actin and means ± SEM in 3 independent experiments are shown with values in control cells set at “1.” Total cell numbers were evaluated at day 3 of differentiation and are presented relative to shCTRL conditions (right, n=6). (**D**) CD36 (top) and CD71 (bottom) were evaluated in shCTRL- and shRPL11-transduced progenitors at day 3 of EPO stimulation. Representative histograms are shown (left) and quantification of positive cells were compared relative to shCTRL cells (right, n=10). (**F**) SRC levels ± SEM are presented for shCTRL- and shRPL11-transduced progenitors (n=3 independent experiments). *p<0.05; ****p<0.0001; ns, not significant

